# NRF2-dependent metabolic reprogramming is required for tumor recurrence following oncogene inhibition

**DOI:** 10.1101/513994

**Authors:** Douglas B. Fox, Ryan Lupo, Laura C. Noteware, Rachel Newcomb, Juan Liu, Jason W. Locasale, Matthew D. Hirschey, James V. Alvarez

## Abstract

Oncogenic signaling pathways both directly and indirectly regulate anabolic metabolism, and this is required for tumor growth. Targeted therapies that inhibit oncogenic signaling have dramatic impacts on cellular metabolism. However, it is not known whether the acquisition of resistance to these therapies is associated with – or driven by – alterations in cellular metabolism. To address this, we used a conditional mouse model of Her2-driven breast cancer to study metabolic adaptations following Her2 inhibition, during residual disease, and after tumor recurrence. We found that Her2 downregulation caused widespread changes in cellular metabolism, culminating in oxidative stress. Tumor cells adapted to this metabolic stress by upregulation of the antioxidant transcription factor, NRF2. Constitutive NRF2 expression persisted during residual disease and tumor recurrence, and NRF2 was both sufficient to promote tumor recurrence, and necessary for recurrent tumor growth. These results are supported by clinical data showing that the NRF2 transcriptional program is activated in recurrent breast tumors, and that NRF2 is associated with poor prognosis in patients with breast cancer. Mechanistically, NRF2 signaling in recurrent tumors induced metabolic reprogramming to re-establish redox homeostasis and upregulate *de novo* nucleotide synthesis. Finally, this NRF2-driven metabolic state rendered recurrent tumor cells sensitive to glutaminase inhibition, suggesting that NRF2-high recurrent tumors can be therapeutically targeted. Together, these data provide evidence that NRF2-driven metabolic reprogramming is required for breast cancer recurrence following oncogene inhibition.

**Significance:** Although tumor recurrence is the leading cause of mortality in breast cancer, the cellular properties that allow tumor cells to evade therapy and form recurrent tumors remain largely uncharacterized. Similarly, very little is known about how tumor metabolism changes following therapy, or whether alterations in cellular metabolism drive tumor recurrence. In this study, we identify the antioxidant transcription factor NRF2 as a critical positive regulator of breast cancer recurrence. We find that NRF2-dependent metabolic reprogramming is both sufficient and required to promote tumor recurrence. Additionally, we demonstrate that the NRF2-driven metabolic state renders recurrent tumors sensitive to glutaminase inhibitors, suggesting a novel therapeutic approach for the treatment of recurrent breast cancer.

## Introduction

Tumor formation requires profound changes in cellular metabolism. These changes are mediated, in part, through the direct regulation of metabolic pathways by oncogenes and tumor suppressors (Boroughs and DeBerardinis, 2015; Pavlova and Thompson, 2016). For example, the PI3K-Akt pathway promotes glucose utilization via several mechanisms, including promoting the membrane localization of glucose transporters (Elstrom et al., 2004). MYC induces transcription of glutaminase and glutamine transporters to induce glutamine utilization (Wise et al., 2008). Her2/neu signaling, in addition to regulating glucose metabolism through the PI3K-Akt pathway, induces high rates of de novo fatty acid synthesis by regulating fatty acid synthase (Menendez et al., 2004) and lipid storage pathways (Wang et al., 2013). These oncogenic pathways are necessary to sustain anabolic metabolism. Metabolic reprogramming is an early event in transformation, and it is required for tumorigenesis (Hanahan and Weinberg, 2011; Tennant et al., 2009).

Targeted therapies that directly inhibit oncogenic signaling have become mainstays in clinical oncology. These drugs can induce dramatic clinical responses (Flaherty et al., 2010; MacConaill et al., 2011), though resistance to targeted therapies is a common problem that limits their efficacy (Lackner et al., 2012; Niederst and Engelman, 2013). While the function of metabolic reprogramming in the development of cancer has been intensively studied, relatively little is known about how therapy-induced oncogene inhibition alters cellular metabolism. Similarly, it is not known whether the acquisition of resistance to targeted therapies is associated with – or driven by – alterations in cellular metabolism.

Several recent studies have begun to address these issues. For instance, it has been shown that inhibition of oncogenic signaling causes an immediate drop in glucose uptake both *in vitro* (Viale et al., 2014; Zhao et al., 2011) and *in vivo* (Alvarez et al., 2014). In response to decreased glucose metabolism, cells that survive the loss of oncogenic signaling upregulate fatty acid oxidation (FAO) pathways (Havas et al., 2017; Viale et al., 2014). Additionally, altered metabolism following oncogene inhibition results in increased levels of reactive oxygen species (ROS) (Havas et al., 2017; Krall et al., 2017; Viale et al., 2014), which contribute to cell death following therapy. These results suggest that tumor cells that survive therapy and eventually develop resistance must adapt to overcome therapy-induced metabolic stress. However, the pathways that promote metabolic adaptations following therapy remain unknown.

To gain insight into how tumor cell metabolism changes following oncogene inhibition, and how these changes contribute to the survival and recurrence of residual cells following therapy, we used a transgenic mouse model of Her2-driven breast cancer (Alvarez et al., 2013; Moody et al., 2005; Moody et al., 2002). In this model, doxycycline (dox) administration to bitransgenic MMTV-rtTA;TetO-Her2 (MTB/TAN) mice leads to Her2 expression and the formation of invasive mammary adenocarcinomas. Removal of doxycycline causes Her2 downregulation and induces tumor regression. However, a small population of cells persists in a dormant state before eventually re-initiating proliferation to form a recurrent tumor. Using this model, we interrogated the metabolic changes that accompany Her2 downregulation, residual disease, and tumor recurrence. We identified the NRF2 pathway as a critical mediator of metabolic reprogramming following oncogene inhibition. NRF2 promotes tumor recurrence by promoting redox homeostasis and nucleotide metabolism in recurrent tumors. As a consequence of this metabolic reprogramming, recurrent tumors with high NRF2 are sensitive to glutaminase inhibition, suggesting a therapeutic strategy for treating recurrent tumors.

## Results

### Her2 Downregulation Induces Metabolic Changes and Loss of Redox Homeostasis

To examine the metabolic consequences of Her2 downregulation, we used an *in vitro* model in which tumor cells maintain their dependency upon Her2. Primary Her2-driven tumors were digested and cultured as mammospheres in non-adherent, serum-free conditions in the presence of dox. Removal of dox from these cells led to a rapid decrease in Her2 levels and in downstream components of the MAPK and Akt-mTOR pathways (Figure S1A and B). This was associated with a loss of proliferation that was evident as early as 2 days (Figure 1A), and an induction of apoptosis starting at 7 days following dox withdrawal (Figure 1B and C). This apoptosis was transient, and by 13 days following dox withdrawal we identified surviving cells that were quiescent but non-apoptotic (Figure 1D). These surviving cells were viable, because re-addition of dox to these mammospheres led to re-induction of proliferation (Figure 1E). Taken together, these results indicate that these mammospheres faithfully model the behavior of tumors in vivo: tumor cells remain dependent upon Her2 for their growth and survival, but a population of cells can survive Her2 downregulation and persist in a viable, non-proliferative state.

**Figure 1:**
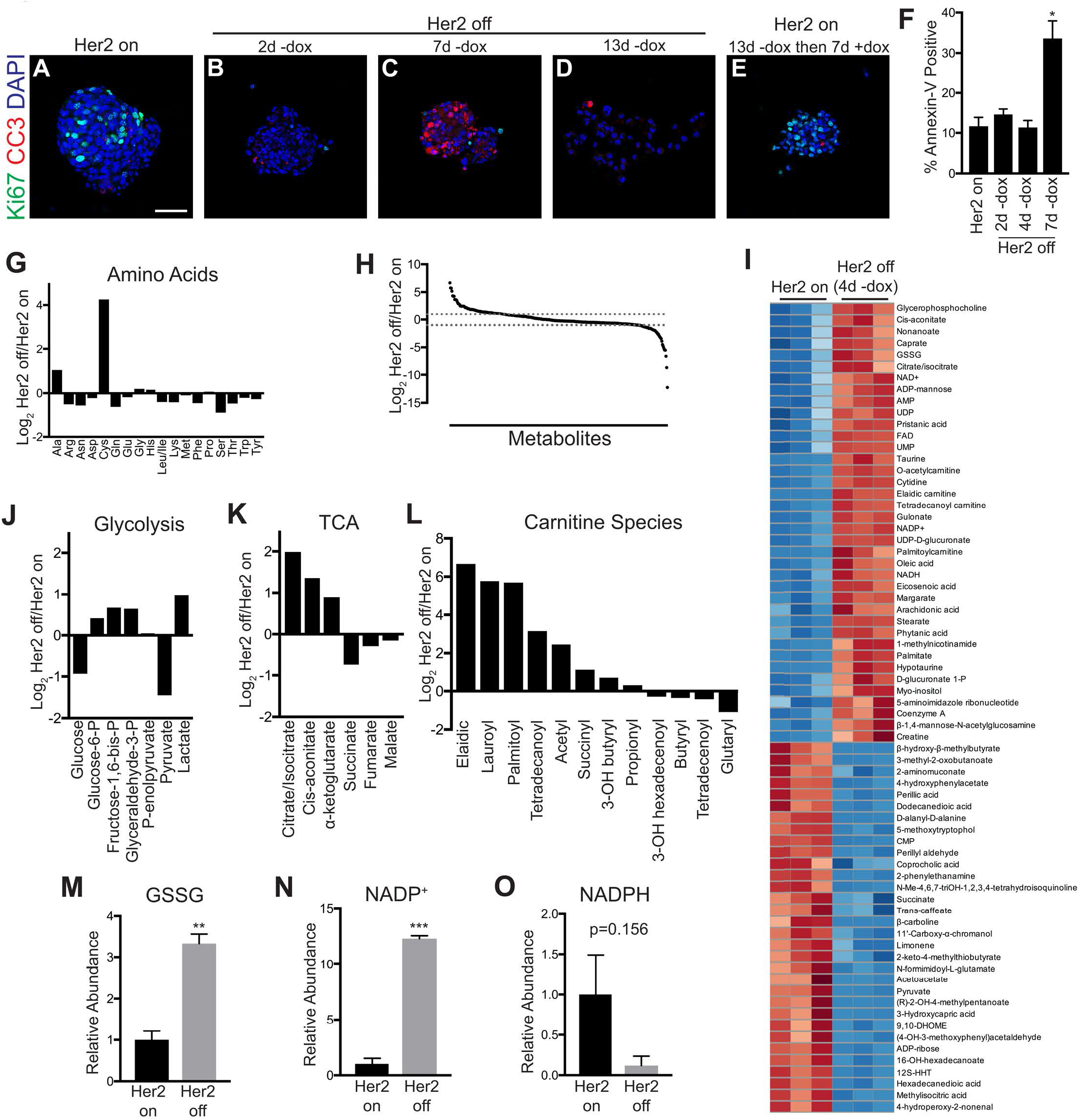
Her2 Downregulation Induces Metabolic Changes. (A-E) Immunofluorescence staining for Ki67 (green) and cleaved caspase-3 (red) in mammospheres cultured in the presence of dox (A; Her2 on), without dox (Her2 off) for 2 days (B), 7 days (C), and 13 days (D), and without dox for 13 days before re-addition of dox for 7 days (E). Scale bar, 50 μm. (F) Percentage of Annexin V-positive cells in mammospheres cultured in the presence of dox (Her2 on), or without dox (Her2 off) for 2, 4, and 7 days. Significance was determined by one-way ANOVA with Tukey’s multiple comparisons test. (n=2) (G) Ratio of metabolite levels between Her2 off and Her2 conditions. Dashed lines represent 2-fold change. (H) Changes in amino acid levels between Her2 off and Her2 conditions. (I) Heatmap showing top 70 altered metabolites following Her2 downregulation in mammospheres. (J-L) Changes in glycolytic intermediates (J), citric acid cycle (TCA) intermediates (K), and carnitine species (L) between Her2 off and Her2 on conditions. (M-O) Relative metabolite levels of oxidized glutathione (GSSG) (M), oxidized NADP+ (N), and reduced NADPH (O) measured in mammospheres cultured in the presence of dox (Her2 on) or without dox (Her2 off). Significance was determined by Student’s t test (n=3). Error bars denote mean ± SEM. *p < 0.05; **p < 0.01; ***p < 0.001. See also Figure S1.

We next used this model to characterize the metabolic changes that occur following Her2 downregulation. To avoid any confounding effects of apoptosis on cell metabolism, we chose a 4-day time-point, which is after the loss of Her2 signaling but prior to the onset of apoptosis (Figure 1F). We used liquid chromatography-mass spectrometry to examine changes in metabolite levels in mammospheres 4 days following Her2 downregulation. The majority of metabolites, including most amino acids (Figure 1G), did not change following Her2 downregulation. Of the 317 measurable metabolites, 70 were significantly increased (fold-change >2, p<0.05) and 26 were significantly decreased (fold-change <0.5, p<0.05) (Figure 1H and I). Glucose and pyruvate levels decreased following Her2 downregulation, while lactate levels increased (Figure 1J). Consistent with this result, we observed an increase in the levels of phosphorylated pyruvate dehydrogenase (p-PDH) (Figure S1C), suggesting that Her2 downregulation diverts pyruvate away from citric acid cycle anaplerosis and toward conversion to lactate. We also observed increased steady-state levels of several citric acid cycle intermediates, including citrate/isocitrate, cis-aconitate, and α-ketoglutarate (Figure 1K). Further, Her2 downregulation induced a considerable increase in the steady-state levels of many acylcarnitines (Figure 1L). Because acylcarnitines are intermediates in fatty acid oxidation, this suggests that these cells have increased levels of fatty acid oxidation (FAO), which is in agreement with findings in similar systems (Havas et al., 2017; Viale et al., 2014).

Interestingly, Her2 downregulation also induced profound changes in the steady-state levels of components of glutathione metabolism. There was a substantial increase in the oxidized form of glutathione (GSSG) following Her2 downregulation (Figure 1M), and this coincided with an increase in NADP+ (Figure 1N) and a decrease in NADPH (Figure 1O). Glutathione is a critical intracellular antioxidant, and it is maintained in its reduced form using reducing capacity of NADPH. Thus, the observed shifts in GSSG, NADP+, and NADPH indicate that Her2 downregulation caused a loss of cellular redox homeostasis. Together, these changes are consistent with our observed changes in lower glycolysis and FAO intermediates, and they suggest that Her2 downregulation causes cells to use FAO for energy generation, which is concomitant with changes in redox homeostasis.

### Her2 Inhibition Increases ROS Levels and Leads to ROS-dependent Cell Death

Changes in the redox states of glutathione and NADP(H) are often indicative of oxidative stress caused by accumulation of reactive oxygen species (ROS). We therefore directly measured the levels of cellular ROS using 2’,7’–dichlorofluorescin diacetate (DCFDA) in mammospheres cultured with or without dox for four days. Her2 downregulation led to a substantial increase in ROS levels (Figure 2A). To ensure that increased ROS was not a secondary consequence of apoptosis, we measured ROS levels two days following dox withdrawal, a timepoint that precedes apoptosis in this model (see Figures 1E and F), and found increased ROS at this early timepoint as well (Figure S2A). To expand these results, we tested the effects of Her2 inhibition in two well-characterized human Her2-amplified breast cancer cell lines, BT474 and SKBR3 cells. Treatment with lapatinib, a small-molecule dual Her2/EGFR inhibitor, led to a dose-dependent increase in ROS levels in both cell lines (Figure 2B and C). We next assessed changes in mitochondrial ROS levels following Her2 inhibition using MitoSox, which specifically measures ROS levels in mitochondria. Mitochondrial ROS levels increased following Her2 downregulation in mammospheres (Figure 2D) as well as in BT474 and SKBR3 cells following lapatinib treatment (Figure 2E and F). Given the well-established role of ROS in inducing cell death, we next asked whether the increased oxidative stress we observed contributes to cell death following Her2 inhibition. Treatment with the antioxidant N-acetylcysteine (NAC) prevented ROS accumulation (Figure 2G) and rescued cell viability following lapatinib treatment (Figure 2H-J and S2B). To further test that this effect was mediated by the antioxidant capacity of NAC, we also treated cells with exogenous GSH, which similarly rescued cell viability following lapatinib treatment (Figure 2K).

**Figure 2:**
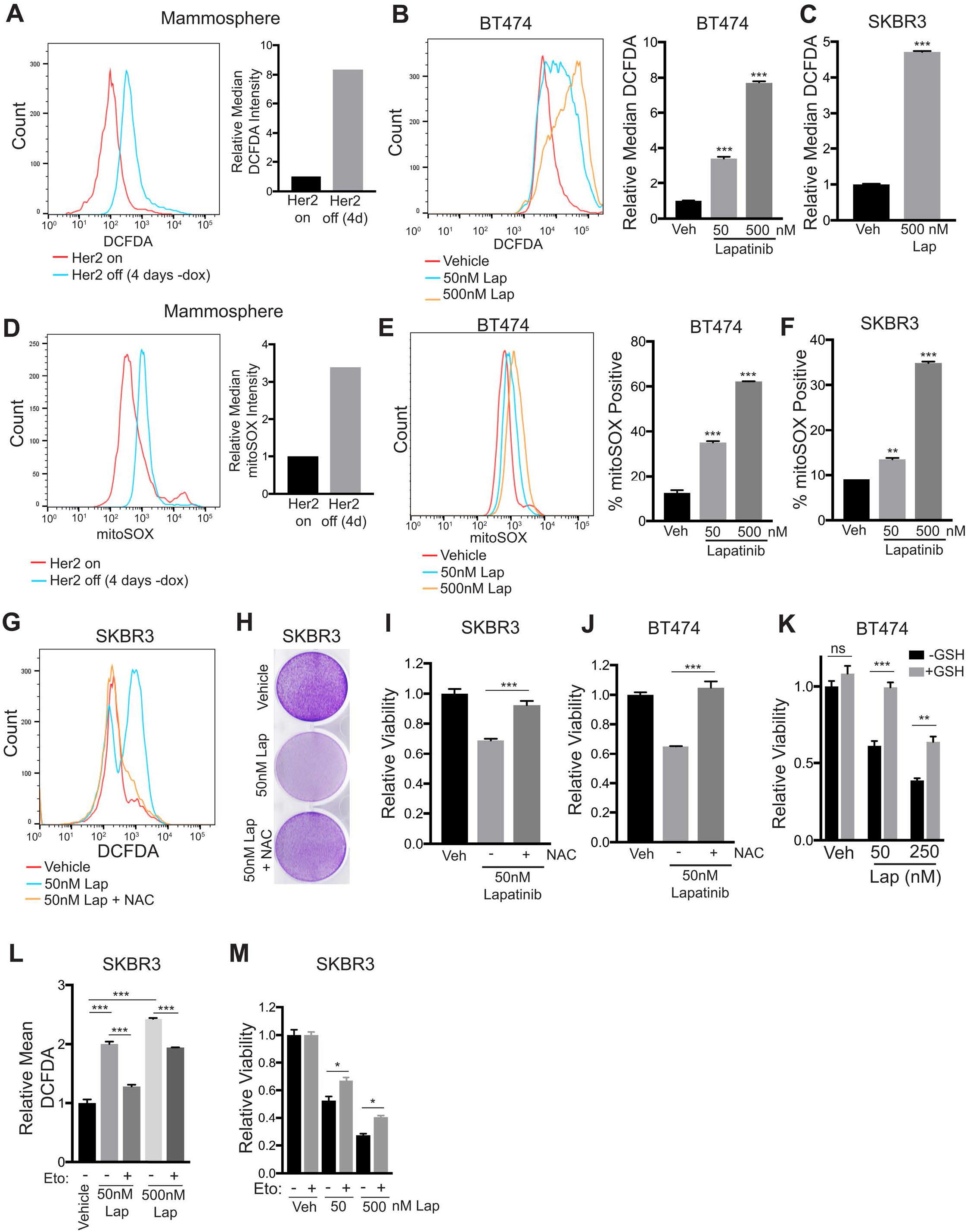
Her2 Inhibition Increases ROS Levels and Leads to ROS-dependent Cell Death. (A) DCFDA staining showing ROS levels in mammospheres cultured with dox or without dox for 4 days. Data are representative of 3 independent experiments. (B-C) DCFDA staining showing ROS levels in BT474 cells (B) and SKRB3 cells (C) treated with lapatinib for 48 hours. Significance was determined by one-way ANOVA (n=2, Tukey’s multiple comparisons test). (D) MitoSOX staining showing mitochondrial superoxide levels in mammospheres cultured with dox or without dox for 4 days. (E-F) MitoSOX staining showing mitochondrial superoxide levels in BT474 cells (E) and SKRB3 cells (F) treated with lapatinib for 48 hours. Significance was determined by one-way ANOVA (n=2, Tukey’s multiple comparisons test). (G) DCFDA staining showing ROS levels in SKBR3 cells treated with lapatinib and 5mM N-acetyl cysteine (NAC). (H) Crystal violet staining showing viability after treatment of SKBR3 cells with lapatinib and 5mM NAC for 6 days. (I-J) Relative viability of SKBR3 cells (I) and BT474 cells (J) treated with lapatinib and 5 mM NAC for 72 hours. Significance was determined by two-way ANOVA (n=3, Tukey’s multiple comparisons test). (K) Relative viability of BT474 cells treated with lapatinib and 10 mM glutathione (GSH). Significance was determined by two-way ANOVA (n=3, Tukey’s multiple comparisons test). (L) DCFDA staining showing ROS levels in SKBR3 cells treated with lapatinib and 100 μM etomoxir (eto). Significance was determined by one-way ANOVA (n=2, Tukey’s multiple comparisons test). (M) Relative viability of SKBR3 cells treated with lapatinib and 100 μM etomoxir (eto) for 72 hours. Significance was determined by two-way ANOVA (n=3, Tukey’s multiple comparisons test). Error bars denote mean ± SEM. *p < 0.05; **p < 0.01; ***p < 0.001. See also Figure S2.

We next wanted to determine the metabolic basis for increased ROS following Her2 inhibition. Recent evidence has shown that inhibition of the glucose dependent oxidative pentose phosphate pathway (PPP), which generates NADPH, is sufficient to induce ROS and cell death in tumor cells (Wang et al., 2017). Additionally, the induction of FAO in cells that survive oncogene downregulation has been shown to cause ROS accumulation (Havas et al., 2017; Viale et al., 2014). Interestingly, we found that treatment with two independent inhibitors of the PPP, dehydroepiandrosterone (DHEA) and 6-aminonicotinamide (6-An), was not sufficient to increase ROS levels in breast cancer cell lines (Figure S2C and D) or mammospheres (Figure S2E). Further, PPP inhibition did not exacerbate induce ROS in combination with lapatinib (Figure S2D). On the other hand, we found that treatment of cells with the FAO inhibitor, etomoxir, prevented ROS accumulation following Her2 inhibition (Figure 2L) and partially rescued cell viability (Figure 2M and S2F). This suggests that Her2 inhibition leads to the induction of FAO, consistent with our observation that acylcarnitines and mitochondrial ROS are enriched following Her2 downregulation (Figure 1K). To explore this further, we tested whether Her2 inhibition induced expression of genes required for FAO. In two independent mammosphere cultures, Her2 downregulation resulted in significant increases in the mitochondrial acylcarnitine transporters, Cpt1a and Cpt1b, and the fatty acid transporter, CD36 (Figure S2G). Additionally, staining of mammospheres with the neutral lipid marker, boron-dipyrromethene (BODIPY), showed that mammospheres contained abundant lipid droplets (Figure S2H). Taken together, these results suggest that the induction of FAO following Her2 inhibition is responsible for ROS accumulation.

### The NRF2 Antioxidant Transcriptional Program Is Activated in Dormant and Recurrent Tumors

The finding that Her2 inhibition leads to ROS-dependent cell death suggests that dormant tumor cells that are able to survive Her2 downregulation may have activated intrinsic cellular antioxidant pathways. The transcription factor NRF2 (nuclear factor (erythroid-derived 2)-like 2, or Nfe2l2) is a critical mediator of the cellular adaptive antioxidant response. NRF2 becomes stabilized in response to oxidative stress and activates genes involved in restoring redox homeostasis (Nguyen et al., 2009). NRF2 has been established as a driver of tumor progression (DeNicola et al., 2011), resistance to therapy (Jiang et al., 2010; Krall et al., 2017; Oshimori et al., 2015), and metastasis (Wang et al., 2016). We first assessed whether NRF2 is stabilized following Her2 downregulation *in vivo*. Western blotting of tumor samples from MTB/TAN mice with primary tumors or shortly after dox withdrawal showed that NRF2 protein levels increase following Her2 downregulation (Figure 3A). Additionally, qRT-PCR analysis showed that the NRF2 target genes, Gclm, Nqo1, and Hmox1, were increased in these tumors (Figure S3A). Similar results were obtained using immunofluorescence staining for NRF2. NRF2 was expressed at low or undetectable levels in primary tumors, but was stabilized and located in the nucleus of tumor cells following Her2 downregulation (Figure 3B). These results indicate that Her2 downregulation leads to NRF2 stabilization *in vivo*, likely as a result of increased ROS.

**Figure 3:**
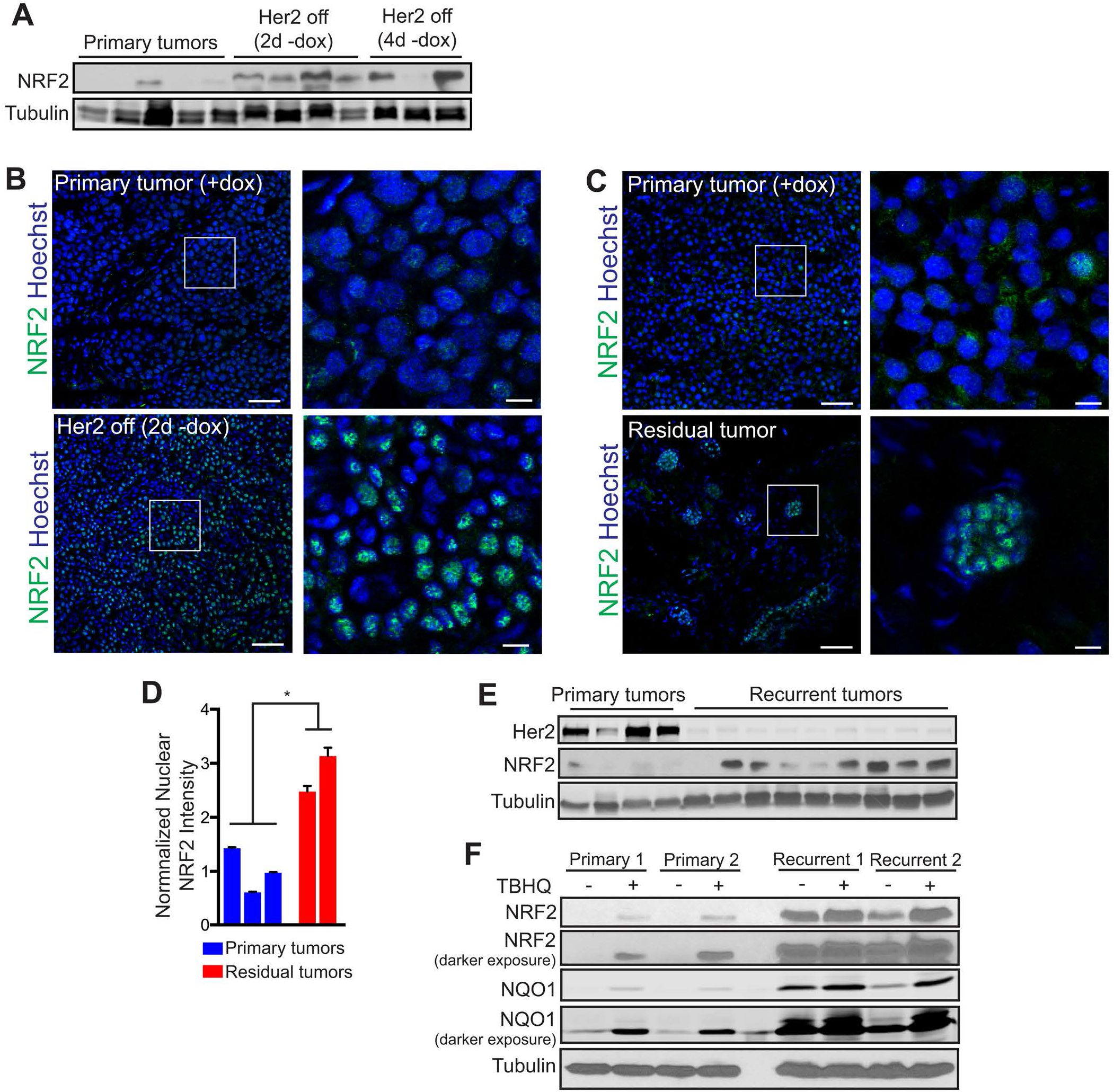
The NRF2 Antioxidant Transcriptional Program is Activated in Dormant and Recurrent Tumors. (A) Western blot for NRF2 in primary tumors and tumors 48 and 96 hours after dox withdrawal (Her2 off). (B-C) Immunofluorescence staining for NRF2 in primary tumors, tumors 48 hours after dox withdrawal (Her2 off) (B) and 5 months after dox withdrawal (residual tumor) (C). Scale bars represent 50 μm (left) and 10 μM (right). White box on left panels indicates magnified section shown in right panels. (D) Quantification of nuclear NRF2 staining from (C) comparing primary tumors and tumors 5 months after dox withdrawal (residual tumors). Significance was determined by Student’s t test of the average values for each group. (E) Western blots for Her2 and NRF2 in primary (n=4) and recurrent (n=9) tumors. (F) Western blots for NRF2 and NQO1 in two independent primary tumor cell lines and two independent recurrent tumor cell lines with or without treatment with the electrophile TBHQ (25 μM). Error bars denote mean ± SEM. ***p < 0.001. See also Figure S3.

A small population of tumor cells survives Her2 downregulation and persists in a dormant state prior to seeding recurrent tumors (Feng et al., 2014). We next wanted to determine if NRF2 expression remains elevated in these residual tumor cells. Immunofluorescence staining for NRF2 in dormant tumors revealed that residual tumors exhibited high nuclear NRF2 staining, suggesting that this antioxidant response persisted in dormant tumors (Figure 3C and D). Similar results were obtained in tumors generated from a lower dose of dox (0.1 mg/ml), which yields more focal residual tumors (Figure S3B).

Residual tumor cells persist for several months before spontaneously re-initiating proliferation to give rise to recurrent tumors. We asked whether these recurrent tumors maintain elevated NRF2. Western blotting showed that the majority of recurrent tumors had high NRF2 levels (Figure 3E). We next compared NRF2 levels in cells derived from independent primary and recurrent tumors. NRF2 levels were elevated in recurrent tumor cells as compared to primary cells (Figure 3F), mimicking the behavior of primary and recurrent tumors *in vivo*. Similarly, NRF2 target genes were elevated in recurrent tumor cells cultured *in vitro* (Figure S3C). Taken together, these results demonstrate that tumors upregulate the NRF2 antioxidant pathway in response to metabolic changes caused by Her2 downregulation, and NRF2 activation persists during dormancy and in recurrent tumors.

### Constitutive NRF2 Activity Promotes Tumor Recurrence

In light of our observation that NRF2 is frequently activated in cancer cells in response to the metabolic stress induced by Her2 inhibition, and remains elevated in residual and recurrent tumors, we next sought to determine if NRF2 activity promotes tumor recurrence. To this end, we ectopically expressed a constitutively active NRF2 (caNRF2) – which lacks the N-terminal domain that binds its negative regulator, KEAP1 (Kobayashi et al., 2002; Kohler et al., 2014) (Figure 4A) – in primary tumor cells. Western blotting and qRT-PCR analysis showed that caNRF2 was expressed in primary tumor cells and was transcriptionally active, evidenced by a marked increase in expression of its target genes Nqo1, Slc7a11, and Gclm (Figure 4B and C). Additionally, basal ROS levels were modestly decreased in cells expressing caNRF2 (Figure S4A). Cohorts of recipient mice on dox were then injected with primary tumor cells expressing either empty vector or caNRF2. Expression of caNRF2 had no effect on the rate of primary tumor formation (Figure S4B). Following primary tumor formation, mice were then removed from dox to induce Her2 downregulation and tumor regression. Both control and caNRF2 tumors fully regressed following dox withdrawal (data not shown). Mice with residual tumors were palpated weekly to monitor the emergence of recurrent tumors. Tumors expressing caNRF2 recurred nearly 3 weeks sooner than control tumors (Figure 4D). Importantly, caNRF2 and its target, NQO1, remained highly expressed throughout the time course of recurrence (Figure S4C). These results indicate that NRF2 activity promotes the formation of recurrent tumors.

**Figure 4:**
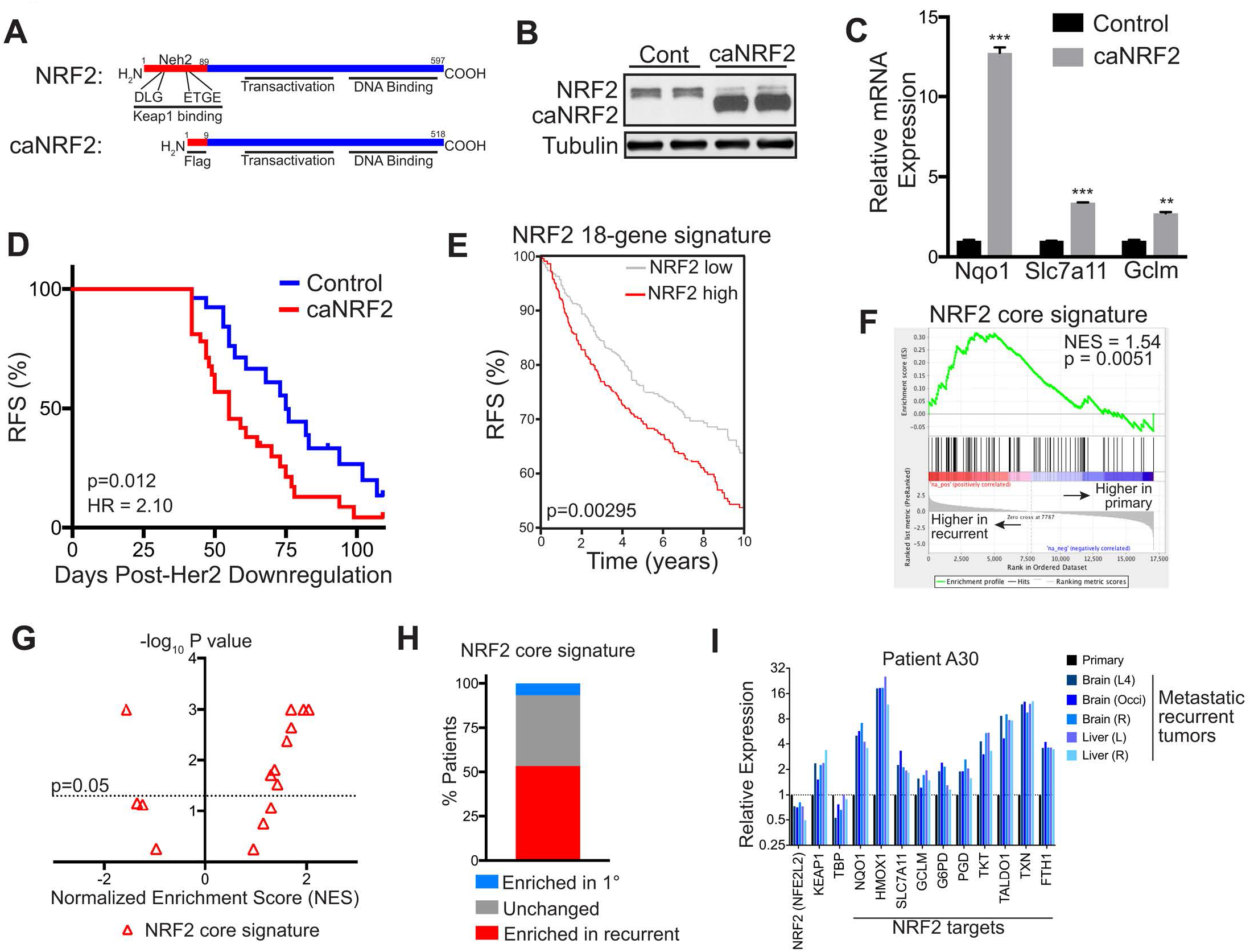
Constitutive NRF2 Activity Promotes Tumor Recurrence. (A) Schematic showing wild-type (top) and constitutively active NRF2 (caNRF2, bottom) proteins. caNRF2 lacks the first 86 amino acids containing the KEAP1 binding domains, DLG and ETGE. (B) Western blot for NRF2 in primary tumor cells expressing and empty vector (Cont) or caNRF2. (C) qPCR analysis of Nqo1, Slc7a11, and Gclm expression in primary tumor cells expressing and empty vector (Control) or caNRF2. Significance was determined by two-way ANOVA with Tukey’s multiple comparisons test (n=2). (D) Kaplan-Meier plot showing recurrence-free survival (RFS) for mice bearing control or caNRF2 tumors. p-value was determined by log-rank (Mantel-Cox) test. (E) Kaplan-Meier plot showing recurrence-free survival (RFS) for breast cancer patients whose tumors have low (gray) or high (red) expression of the NRF2 18-gene signature. (F) Gene set enrichment analysis of matched primary and recurrent breast tumors showing enrichment of the NRF2 core signature in recurrent tumors. (G) Volcano plot showing enrichment scores for the NRF2 core signature in recurrent tumors in individual patients. (H) The percent of patients whose recurrent tumors had a significantly enriched (red), unchanged (grey), or significantly decreased (blue) NRF2 core signature. (I) Expression of NRF2, KEAP1, TBP, and 10 canonical NRF2 target genes in the primary tumor and matched recurrent tumors from a single patient. Error bars denote mean ± SEM. **p < 0.01; ***p < 0.001. See also Figure S4.

We next wanted to determine if NRF2 expression was correlated with poor clinical prognosis in breast cancer patients. We used a recently curated NRF2 gene signature (Romero et al., 2017) (NRF2 core signature), consisting of 108 genes, and a smaller gene set consisting of 18 canonical target genes (NRF2 18-gene signature). High expression of both gene sets was correlated with an increased risk of recurrence in a cohort of over 1800 breast cancer patients (Ringner et al., 2011) (Figure 4E and S4D). These results suggest that high NRF2 activity in primary tumors may promote recurrence in breast cancer. We next wanted to determine if there was direct evidence for NRF2 activation in recurrent breast cancers in humans. For this, we used RNA sequencing data from a rapid autopsy study that collected matched primary and metastatic recurrent tumors (Siegel et al., 2018). We first ranked genes according to their fold-change between primary and recurrent tumors in each patient, averaged the fold-change in expression across all patients, and then performed gene set enrichment analysis (GSEA) to identify gene expression changes in recurrent tumors. GSEA showed that the NRF2 core gene signature was significantly enriched in recurrent tumors (Figure 4F). Notably, this analysis also showed that the “Reactive Oxygen Species Pathway” and “Fatty Acid Metabolism” hallmark gene signatures were enriched in recurrent tumors (Figure S4E). We then repeated GSEA in individual patients to determine the frequency of NRF2 enrichment in recurrent tumors. GSEA showed that the NRF2 core signature was significantly enriched in recurrent tumors in 8 patients (53.3%), not significantly changed in 6 patients (40%), and significantly decreased in only 1 patient (6.7%) (Figure 4G and H). To further illustrate this, we focused on the expression of canonical NRF2 target genes in 4 individual patients whose recurrent tumors had enrichment for a NRF2 signature. The expression of NRF2 itself (NFE2l2), KEAP1, and a housekeeping gene, TBP, was not significantly altered in patients with high NRF2 enrichment scores. However, NRF2 target genes were consistently upregulated in recurrent tumors from these patients as compared to matched primary tumors (Figure 4I and S4F). Together, these results show that NRF2 activity is frequently increased in recurrent breast cancer in humans.

### NRF2 is Required for Recurrent Tumor Growth In Vivo

Having established that expression of constitutively active NRF2 is sufficient to promote tumor recurrence, we next asked whether NRF2 activation is necessary for the growth of recurrent tumors. To do this, we knocked down NRF2 expression in recurrent tumor cells with high NRF2 (see Figure 3F). Expression of independent shRNAs targeting NRF2 suppressed NRF2 mRNA levels in recurrent cells by 75-90% (Figure 5A). Consistent with this, there was a significant reduction in the mRNA expression of NRF2 target genes, Nqo1, Slc7a11, Gclm, and Hmox1 (Figure 5A) and a reduction in the protein levels of NRF2 and NQO1 (Figure 5B). We next assessed the effect of NRF2 knockdown on tumor cell proliferation in vitro and tumor growth *in vivo*. NRF2 knockdown had no effect on the proliferation rate of recurrent tumor cells in vitro, as measured by the rate of population doublings (Figure 5C). However, when implanted into the mammary fat pad of mice, the growth of NRF2-knockdown tumors was significantly impaired (Figure 5D and E). These results show that NRF2 activation is required for the growth of recurrent tumors *in vivo*.

**Figure 5:**
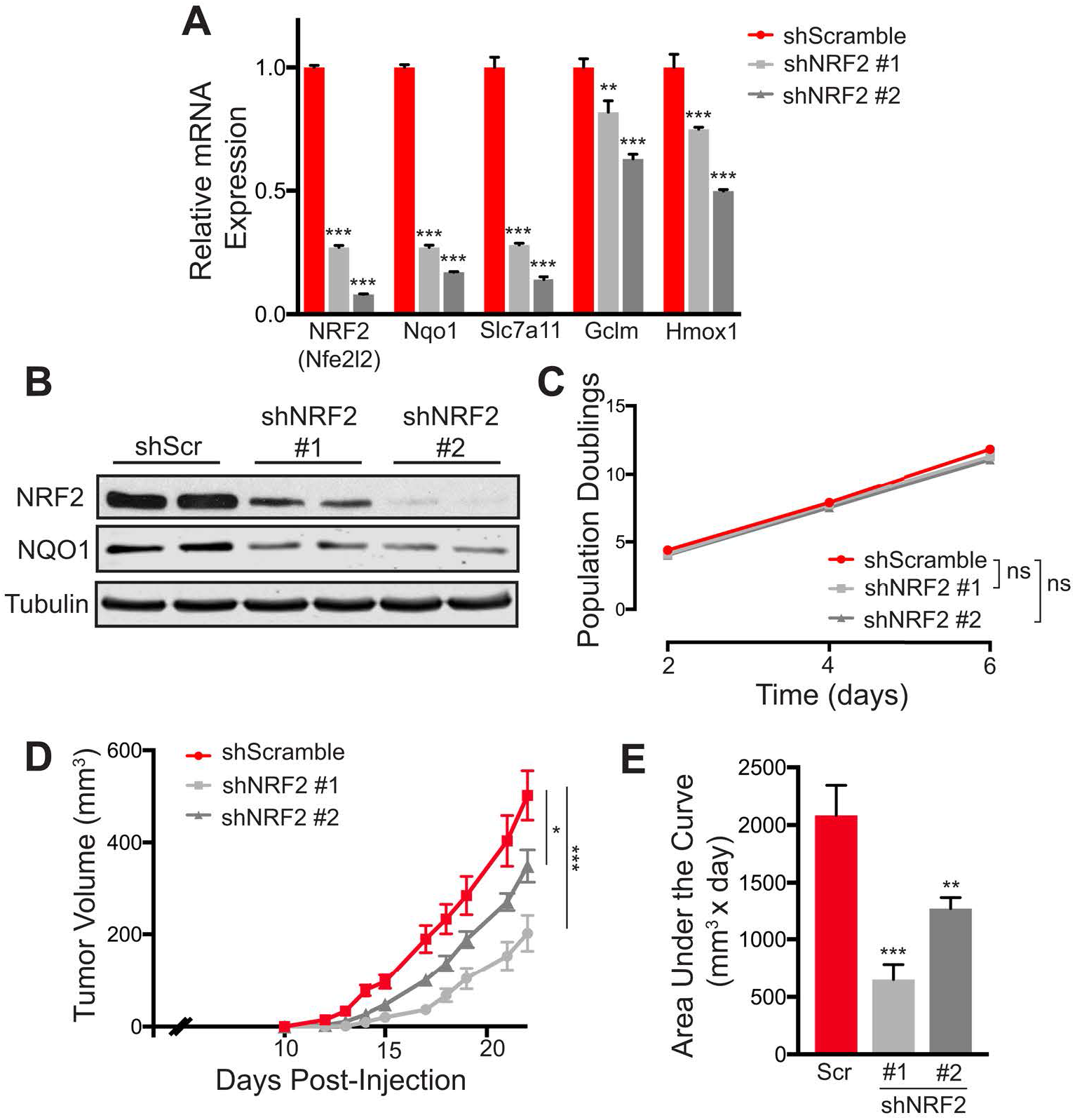
NRF2 is Required for Recurrent Tumor Growth In Vivo. (A) qPCR analysis of NRF2 (Nfe2l2), Nqo1, Slc7a11, Gclm, and Hmox1 expression in recurrent tumor cells expressing scrambled or NRF2-targeting shRNA. Significance was determined by two-way ANOVA with Tukey’s multiple comparisons test (n=2). (B) Western blot for NRF2 and NQO1 in control (shScr) and NRF2-knockdown (shNRF2) recurrent tumor cells. (C) Population doublings in control (shScr) and NRF2-knockdown (shNRF2) recurrent tumor cells. (D-E) Tumor growth curves (D) and area under the curve analysis (E) for tumors generated from control (shScr) and NRF2-knockdown (shNRF2) recurrent tumor cells. Significance was determined by one-way ANOVA using the mean tumor volumes at experiment endpoint (n=16 per condition, Tukey’s multiple comparisons test). Error bars denote mean ± SEM. ns p > 0.05; *p < 0.05; **p < 0.01; ***p < 0.001.

### NRF2 is Activated in Recurrent Tumors Through a Noncanonical Mechanism

We next wanted to determine the mechanism of constitutive NRF2 protein expression in recurrent tumor cells. NRF2 is canonically regulated by KEAP1, which sequesters NRF2 in the cytoplasm and targets it for ubiquitylation by the CUL3 E3 ubiquitin ligase, and it is rapidly degraded by the proteasome under basal conditions (Furukawa and Xiong, 2005). Upon oxidative stress, however, cysteine residues on KEAP1 becomes oxidized, and this induces a conformational change that limits NRF2 ubiquitylation. Therefore, we first wanted to determine if high NRF2 levels in recurrent tumors cells are a consequence of high ROS levels in these cells. We found that basal ROS levels in recurrent tumor cells were either equal to, or lower than, ROS levels in primary tumor cells (Figure 6A). Similarly, mitochondrial ROS levels were not consistently higher in recurrent tumor cells (Figure 6B). Additionally, metabolomic analysis revealed that GSSG levels were equivalent between primary and recurrent cells (Figure 6C). These three independent measurements of ROS levels all indicated that recurrent tumor cells do not exhibit persistent oxidative stress, suggesting that NRF2 activation in these cells is not a consequence of elevated ROS. We next tested whether antioxidant treatment decreased NRF2 levels. Treatment of recurrent tumor cells with NAC reduced ROS levels (Figure S6A), but protein expression of NRF2 and its target, NQO1, were only modestly affected (Figure 6D). Together, these results suggest that NRF2 is activated in recurrent tumor cells independent of ROS.

**Figure 6:**
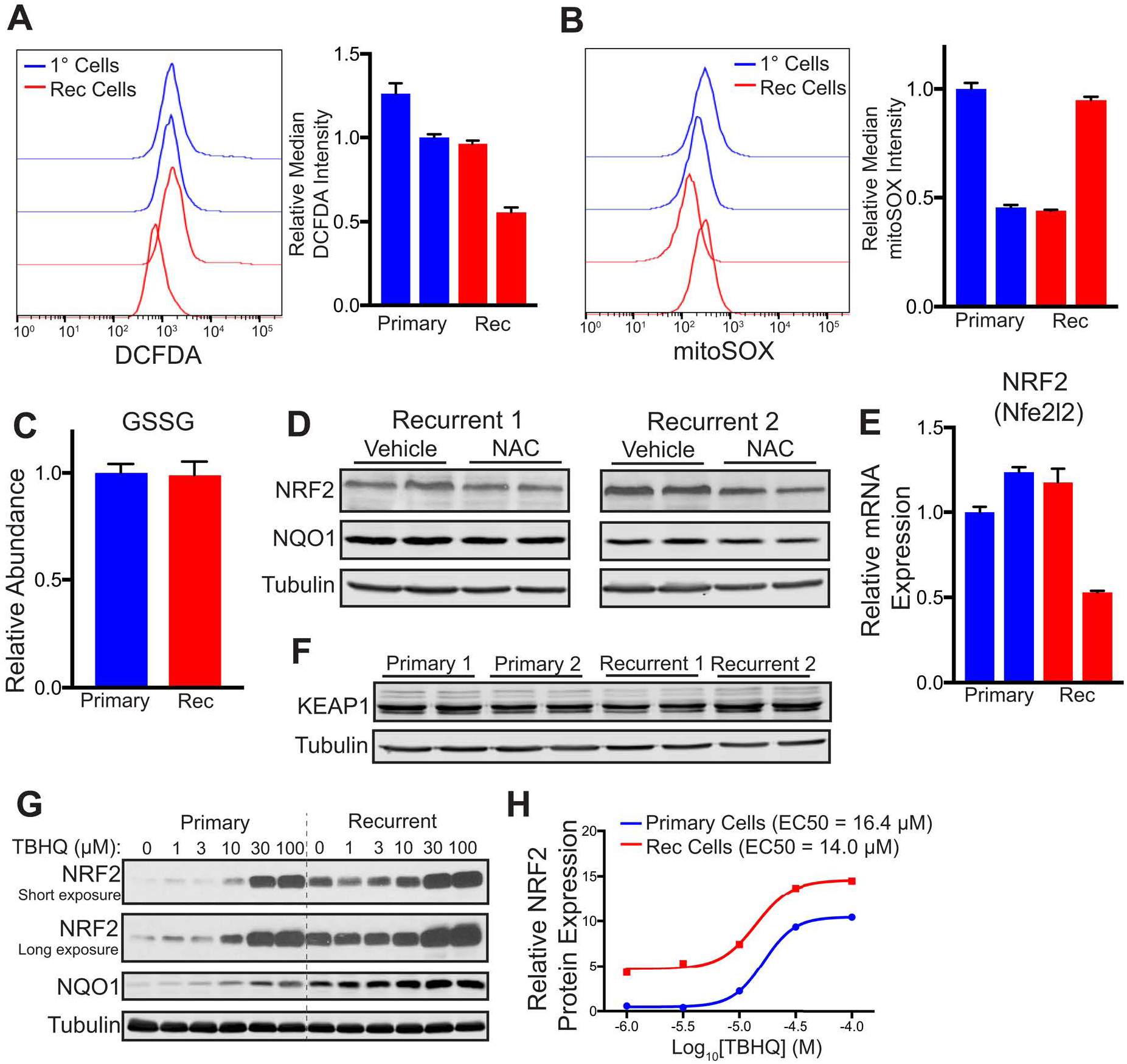
NRF2 Is Non-Canonically Activated in Recurrent Tumor Cells. (A-B) DCFDA staining showing ROS levels (A) and mitoSOX staining showing mitochondrial superoxide levels (B) in two independent primary cell lines and two independent recurrent (Rec) cell lines (n=2 per cell line). (C) Relative levels of oxidized glutathione (GSSG) in a primary cell line and a recurrent cell line (n=3). (D) Western blots for NRF2 and NQO1 in two independent recurrent cell lines treated with 5 mM NAC. (E) qPCR analysis of NRF2 (Nfe2l2) expression in two independent primary cell lines and two independent recurrent (Rec) cell lines (n=2 per cell line). (F) Western blot for KEAP1 in two independent primary cell lines and two independent recurrent (Rec) cell lines. (G) Western blots for NRF2 and NQO1 in a primary cell line and a recurrent cell line treated with the indicated dose of TBHQ for 16 hours. (H) Quantification of NRF2 expression (normalized to Tubulin) from (G). Error bars denote mean ± SEM. See also Figure S5

Several noncanonical mechanisms of NRF2 activation have been reported. These mechanisms include increased transcription of the NRF2 gene (NFE2L2) (Wakabayashi et al., 2014), decreased expression of the NRF2 negative regulator, KEAP1, and somatic mutations of KEAP1 and NRF2 (Hast et al., 2014; Padmanabhan et al., 2006). Both NRF2 mRNA levels and KEAP1 protein levels were not significantly different between primary and recurrent tumor cells (Figure 6E and F). Additionally, neither NRF2 (Nfe2l2) nor Keap 1 was mutated in recurrent tumors (data not shown). We wanted to test if NRF2 remained regulated by KEAP1 in primary and recurrent cells by testing NRF2 stabilization by a KEAP1 oxidizing agent, tert-Butylhydroquinone (TBHQ). Treatment of primary and recurrent cells with increasing doses of TBHQ led to robust increases in NRF2 protein in both cell types (Figure 6G). Analysis of dose-response curves revealed that both cell lines had similar EC_50_ values for TBHQ-induced NRF2 stabilization (Figure 6H), suggesting that NRF2 remains regulated by KEAP1 in recurrent tumor cells, but it has a lower ROS threshold for stabilization. Together, these results indicate that NRF2 is activated in recurrent tumor cells through a noncanonical mechanism.

### NRF2 is Required for Redox Homeostasis in Recurrent Tumor Cells

We next wanted to examine the metabolic basis of the requirement for NRF2 activity in recurrent tumors. We reasoned that NRF2 expression in recurrent tumors may control a similar set of metabolic pathways as Her2 does in primary tumor cells. To gain insight into these pathways, we performed metabolomics on recurrent tumor cells with and without NRF2 knockdown, as well as primary tumor cells grown in the presence or absence of dox (Her2 on or off). We first sought to identify global changes in metabolic pathways between control and NRF2-knockdown cells using pathway enrichment analysis. We found that “Glutathione Metabolism” was one of the top altered pathways following NRF2 knockdown in recurrent tumor cells (Figure 7A). We focused on levels of reduced glutathione (GSH) as a readout of this pathway, and examined GSH levels across all samples. GSH levels decreased following Her2 downregulation in primary tumor cells (Figure 7B). Recurrent tumor cells had reestablished high GSH levels, and in fact GSH were 5-fold higher in recurrent tumor cells as compared to primary tumor cells (Figure 7B). Importantly, NRF2 knockdown in recurrent tumor cells led to a near complete loss of GSH (Figure 7B). Taken together, these results suggest that Her2 signaling regulates GSH levels in primary tumor cells, while NRF2 controls GSH levels in recurrent tumor cells.

**Figure 7:**
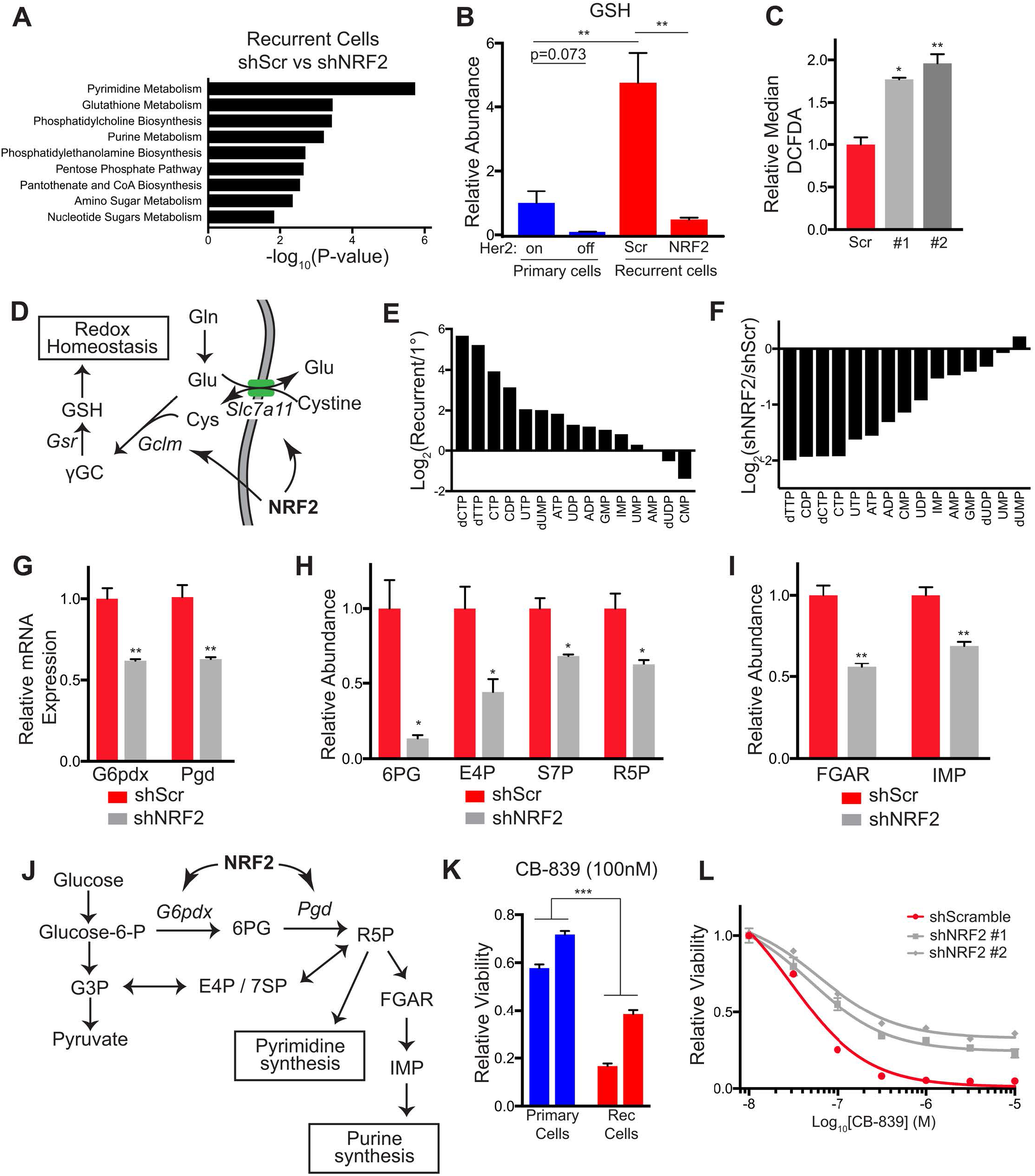
NRF2 Directs Metabolic Reprogramming in Recurrent Tumor Cells. (A) Pathway analysis showing top altered metabolic pathways between control (shScr) and NRF2-knockdown (shNRF2) recurrent tumor cells. (B) Relative levels of reduced glutathione (GSH) in primary tumor cells cultured with dox (Her2 on) and without dox for 7 days (Her2 off), and in control (shScr) and NRF2-knockdown (shNRF2) recurrent tumor cells. Significance was determined by Student’s t test (n=3 per condition). (C) DCFDA staining showing ROS levels in control (shScr) and NRF2-knockdown (shNRF2 #1 and 2) recurrent tumor cells. Significance was determined by one-way ANOVA with Tukey’s multiple comparisons test (n=2). (D) Schematic showing NRF2 regulation of Slc7a11 and Gclm, which are required for glutathione synthesis and promote redox homeostasis. (E) Waterfall plots showing the fold-change in nucleotide levels between primary (1°) and recurrent tumor cells. (F) Waterfall plots showing the fold-change in nucleotide levels between control (shScr) and NRF2-knockdown (shNRF2) recurrent tumor cells. (G) qPCR analysis of G6pdx and Pgd expression in control (shScr) and NRF2-knockdown (shNRF2) recurrent tumor cells. Significance was determined by two-way ANOVA with Tukey’s multiple comparisons test (n=2). (H) Relative levels of the pentose phosphate pathway metabolites, 6-Phosphogluconate (6PG), Erythrose 4-phosphate (E4P), Sedoheptulose 7-phosphate (S7P), and Ribose 5-phosphate (R5P), in control (shScr) and NRF2-knockdown (shNRF2) recurrent tumor cells. Significance was determined by Student’s t test (n=3 per condition). (I) Relative levels of nucleotide precursors, 5'-Phosphoribosyl-N-formylglycineamide (FGAR) and Inosine monophosphate (IMP) in control (shScr) and NRF2-knockdown (shNRF2) recurrent tumor cells. Significance was determined by Student’s t test (n=3 per condition). (J) Schematic showing NRF2 regulation of oxidative pentose phosphate pathway genes (G6pdx and Pgd) and precursors of pyrimidine and purine biosynthesis (6PG, E4P, S7P, R5P, FGAR, and IMP). (K) Relative viability of primary and recurrent cell lines treated with the glutaminase inhibitor, CB-839, normalized to vehicle-treated condition (n=3 per condition). Significance was determined by two-way ANOVA with Tukey’s multiple comparisons test (n=3). p < 0.001 for all pairwise comparisons between primary and recurrent cells. (L) Dose response curves showing relative viability in control (scScr) and NRF2-knockdown (shNRF2) recurrent tumor cells treated with indicated doses of CB-839. Error bars denote mean ± SEM. *p < 0.05; **p < 0.01. ***p < 0.001.

We next examined the mechanistic basis of these changes, focusing on NRF2’s function as a transcription factor. NRF2 knockdown in recurrent tumor cells led to a large reduction in the expression of enzymes involved in de novo glutathione biosynthesis, including the cystine transporter Slc7a11 and a subunit of glutamate-cysteine ligase, Gclm (See figure 5A). Given the decrease in glutathione levels, we speculated that NRF2 knockdown may lead to oxidative stress. Indeed, DCFDA staining showed a 2-fold increase in ROS levels in NRF2 knockdown cells (Figure 7C). These results indicate that NRF2 stabilization is essential for maintenance of redox homeostasis in recurrent tumor cells (Figure 7D).

### NRF2 Directs Metabolic Reprogramming in Recurrent Tumor Cells

We next examined additional metabolic pathways regulated by NRF2 in recurrent tumor cells. Nearly half of the differentially regulated pathways between control and NRF2-knockdown cells related to nucleotide metabolism (“Pyrimidine Metabolism,” “Purine Metabolism,” “Pentose Phosphate Pathway,” and “Nucleotide Sugars Metabolism”) (Figure 7A). Levels of both purine and pyrimidine nucleotides were higher in recurrent tumor cells, which have high NRF2 expression, as compared to primary cells (Figure 7E). Consistent with this, NRF2 knockdown in recurrent tumor cells reduced the levels of these nucleotides (Figure 7F), indicating that NRF2 drives high nucleotide levels in recurrent tumor cells.

The pentose phosphate pathway shunts glucose carbon from glycolysis to form ribose sugars, which are critical for *de novo* nucleotide biosynthesis (Lane and Fan, 2015), and NRF2 has been reported to regulate enzymes of this pathway (Mitsuishi et al., 2012). We next explored whether NRF2 regulation of this pathway is responsible for its role in maintaining nucleotide levels in recurrent cells. qRT-PCR analysis showed that expression of the oxidative PPP enzymes G6pdx and Pgd was decreased following NRF2 knockdown (Figure 7G), but expression of enzymes in the non-oxidative PPP was not substantially altered (Figure S7A). Many metabolite intermediates of the PPP were decreased following NRF2 knockdown (Figure 7H), suggesting that NRF2 directs glucose carbon from glycolysis to form ribose-5-phosphate, which is essential for both purine and pyrimidine synthesis. Finally, we examined intermediates of purine biosynthesis, and found that the metabolites FGAR (5'-Phosphoribosyl-N-formylglycineamide) and IMP (Inosine monophosphate) were also decreased in NRF2 knockdown cells (Figure 7I). Together, these results demonstrate that NRF2-dependent expression of PPP enzymes in recurrent tumor cells results in increased levels of nucleotide precursors and nucleotides. Furthermore, these findings suggest that NRF2 activity is required in recurrent tumors both to maintain redox homeostasis and to drive anabolic pathways that enable tumor growth (Figure 7J).

### Recurrent Tumors with High NRF2 Expression are Sensitive to Glutaminase Inhibition

We next asked if the dependency of recurrent tumor cells for NRF2 activity induces metabolic vulnerabilities that might be targeted to prevent the growth of recurrent tumors. It has recently been shown in non-small cell lung cancer models that constitutive NRF2 activity induced by Keap1 loss confers sensitivity to glutaminase inhibition (Romero et al., 2017; Sayin et al., 2017). We therefore asked whether the elevated NRF2 expression found in recurrent tumor cells renders these cells sensitive to glutaminase inhibition. In agreement with this, the glutaminase inhibitors CB-839 and BPTES inhibited the growth of recurrent tumor cells but not primary tumor cells (Figure 7K and S7B). To test if this sensitivity is mediated by high NRF2 levels, we measured the response of recurrent tumor cells with NRF2 knockdown to glutaminase inhibition. NRF2 knockdown partially rescued cell viability in response to CB-839 and BPTES (Figure 7L and S7C). Interestingly, caNRF2 expression in primary tumor cells was not sufficient to induce sensitivity to glutaminase inhibition (Figure S7D). Together, these results indicate that the NRF2-driven metabolic state renders recurrent breast cancer cells sensitive to glutaminase inhibition.

Glutaminase catalyzes the conversion of glutamine to glutamate, which can be used as an anaplerotic citric acid cycle substrate, for glutathione synthesis, or for secretion by the Xct antiporter in exchange for cystine uptake, which is also used for glutathione synthesis. We wanted to determine if glutaminase activity was required for maintaining redox homeostasis through GSH synthesis in recurrent tumor cells. Interestingly, BPTES treatment resulted in a decrease in ROS levels (Figure S7E), indicating that glutaminase inhibition does not induce oxidative stress. We next asked if replenishing cells with an anaplerotic substrate downstream of glutaminase would rescue viability after glutaminase inhibition. Treatment of cells with dimethyl-α-ketoglutarate (dm-αKG), a cell-permeable form of the TCA intermediate α-ketoglutarate, partially rescued recurrent tumor cell viability following CB-839 treatment (Figure S7F). This is consistent with a recently published model (Sayin et al., 2017), whereby cells with high NRF2 directs glutamate for GSH synthesis and Xct export, thereby limiting the glutamate available for anaplerosis and rendering cells hypersensitive to decreases in glutaminase activity. Together, these results indicate that NRF2 activity, while required for the growth of recurrent tumors, renders them sensitive to glutaminase inhibition.

## Discussion

In this study we examined how tumor cell metabolism changes in response to oncogene inhibition, and elucidated the mechanism of metabolic reprogramming in recurrent tumors. We found that inhibition of Her2 signaling induced substantial and specific changes in cellular metabolism, including changes in metabolites suggestive of a switch from glycolysis to fatty acid oxidation. These metabolic changes resulted in a loss of redox homeostasis and increased ROS, findings consistent with recent reports in similar models (Havas et al., 2017; Viale et al., 2014). We found that treatment with antioxidants could rescue cell death following Her2 inhibition, suggesting that tumor cells that are able to adapt and overcome redox perturbations might be more likely to survive therapy and form recurrent tumors. Consistent with this, we found that tumors responded to Her2 downregulation by activating the antioxidant transcription factor NRF2, which promoted tumor recurrence. We additionally found that a NRF2 transcriptional program is frequently activated in human metastatic recurrent breast cancer, and high expression of NRF2 gene signatures was correlated with poor prognosis, demonstrating the clinical relevance of these metabolic changes. This study highlights an important role for tumor cell adaptation to therapy-induced metabolic stress, and identifies a novel role for NRF2 in promoting breast cancer recurrence by reprogramming cellular metabolism.

Antioxidant pathways are necessary for tumorigenesis (DeNicola et al., 2011; Harris et al., 2015) and tumor progression (Deblois et al., 2016; Takahashi et al., 2018). Recent studies have shown that residual tumor cells experience oxidative stress (Havas et al., 2017; Viale et al., 2014). Consistent with these results, we found that Her2 inhibition led to dysregulation of GSH metabolism and resulted in ROS accumulation and impaired viability. This suggests that re-establishing GSH levels after oncogene inhibition represents a metabolic barrier that tumor cells need to overcome in order to form recurrent tumors. Our results demonstrate that NRF2 activation is a frequent mechanism by which tumor cells accomplish this. As a result, recurrent tumors acquired a dependency on NRF2, which reinforces high levels of GSH metabolism and bolsters their antioxidant defense. This finding is in agreement with several recent studies that have demonstrated the importance of antioxidant pathways for acquired resistance to targeted therapies in breast cancer cell lines (Deblois et al., 2016; Park et al., 2016). Similarly, genetic activation of NRF2 has been recently shown to promote resistance to targeted therapies in lung cancer cells (Krall et al., 2017). These studies support our finding that restoring redox homeostasis through NRF2 activation is imperative for tumor recurrence.

NRF2 can have pleiotropic effects on cell metabolism beyond regulating glutathione metabolism and combating ROS. In particular, NRF2 can activate anabolic pathways, including the nucleotide synthesis (Mitsuishi et al., 2012) and serine synthesis pathways (DeNicola et al., 2015). We found that NRF2 transcriptionally activated genes of the oxidative PPP. This pathway oxidizes glucose-6-phosphate to generate both NADPH, which can be utilized to maintain GSH levels, and ribose sugars required for *de novo* nucleotide synthesis. We found that NRF2 activation in recurrent tumor cells is required to maintain high levels of nucleotides, suggesting that NRF2 orchestrates anabolic metabolism in addition to GSH metabolism in these tumors. We postulate that residual tumor cells with high NRF2 activity may more readily access anabolic pathways, such as nucleotide metabolism, and NRF2 activity during residual disease might potentiate tumor recurrence by priming cells for anabolic metabolism required to resume proliferation.

Our results also demonstrate that the function of NRF2 in mammary tumors is context specific. Primary tumors had low basal levels of NRF2, and ectopic expression of a constitutively active NRF2 did not accelerate tumor growth. Additionally, NRF2 expression in primary tumor cells was not sufficient to induce sensitivity to glutaminase inhibition. In contrast, NRF2 expression accelerated tumor recurrence, and NRF2 knockdown significantly impaired recurrent tumor growth. These results suggest that, in primary tumors, oncogenic Her2 signaling is sufficient to orchestrate antioxidant and anabolic metabolism required for rapid proliferation. This notion is consistent with previous findings that Her2 expression rescues metabolic defects and oxidative stress in mammary acini (Schafer et al., 2009). It is possible that NRF2 and Her2 regulate a convergent set of metabolic pathways, such that NRF2 does not provide an additional benefit to primary tumors and does not impart a dependency on glutaminase. After Her2 downregulation, however, tumor cells become reliant on NRF2 activity to maintain redox homeostasis and *de novo* nucleotide synthesis, and these pathways are indispensable for sustained proliferation.

While KEAP1 and NRF2 mutations are frequently found in lung cancer, they are not common in breast cancers, though recurrent breast cancers have not been extensively sequenced. Interestingly, we found that NRF2 is constitutively activated in recurrent mouse tumors even in the absence of activating genetic mutations. NRF2 activation was not due to increased ROS since recurrent tumors had low basal ROS levels. Further, NRF2 protein levels were increased by the oxidizing agent, TBHQ. This strongly suggests that NRF2 remains regulated by KEAP1, but its threshold for stabilization is much lower in recurrent tumor cells. These observations are consistent with recent reports describing noncanonical modes of NRF2 regulation (Tamir et al., 2016). For instance, post-translational modifications of NRF2 and KEAP1 can inhibit NRF2 degradation in the absence of ROS (Bollong et al., 2018; Chen et al., 2017; Huang et al., 2002). In addition, expression of proteins containing KEAP1-binding domains can compete with NRF2 for KEAP1 binding, resulting in increased basal NRF2 protein expression (Ge et al., 2017; Komatsu et al., 2010; Mulvaney et al., 2016). Further studies will be required to elucidate the mechanism of noncanonical NRF2 stabilization in recurrent breast tumors. A better understanding of these pathways could suggest therapeutic targets for recurrent tumors.

Breast cancer recurrence poses a substantial clinical problem, and therapeutic methods to treat recurrent tumors are currently unavailable. We found that NRF2-driven recurrent tumors acquired a metabolic dependency on GSH metabolism that rendered them sensitive to glutaminase inhibition, suggesting a novel therapeutic approach for the treatment of recurrent breast cancer.

## Experimental Procedures

### Animals

Animal care and animal experiments were performed with the approval of, and in accordance with, guidelines of the Duke University IACUC. Mice were housed under barrier conditions with 12-hour light/12-hour dark cycles. Bitransgenic MTB;TAN mice were generated and tumors were induced as previously described (Alvarez et al., 2013).

### Orthotopic recurrence assays

Orthotopic tumor recurrence assays were performed as described (Alvarez et al., 2013). Cohorts of 14 6-week old immunocompromised mice (nu/nu) under dox administration were injected orthotopically in the 4^th^ inguinal mammary fat pad with 1x10^6^ primary tumor cells (expressing either caNRF2 or an empty vector). Once tumors reached 5 mm (2-3 weeks), dox was be removed to initiate oncogene down-regulation and tumor regression. Mice were be palpated biweekly to monitor tumor recurrence, which was defined as reaching 5x5mm. Differences in recurrence-free survival between control and experimental cohorts were compared using Kaplan-Meier survival curves, and evaluated by the hazard ratio and p-value calculated from a log-rank (Mantel-Cox) test (Alvarez et al., 2013).

### Orthotopic tumor growth assays

Cohorts of 14 6-week old immunocompromised mice (nu/nu) were injected orthotopically in the 4^th^ inguinal mammary fat pad with 1x10^5^ recurrent tumor cells (expressing either a scrambled or NRF2 targeting shRNA). Tumor growth was monitored by palpation, and tumor AUC was calculated using the formula [(vol_1_ + vol_2_)/2]*(day_2_-day_1_).

### Residual tumor samples

Primary tumors were generated in MTB;TAN mice provided with either 2mg/mL or 0.1mg/mL dox water. After primary tumor formation, dox was removed to induce tumor regression. Mammary glands previously bearing primary tumors were harvested 148 (2mg/mL) and 56 (0.1mg/mL) days after dox withdrawal and frozen in OCT.

### Tissue culture and reagents

Mammosphere cultures were generated by digesting primary tumors and plating single cell suspensions on Poly(2-hydroxyethyl methacrylate) (Poly-hema, Sigma) coated plates. Tumor chunks were digested with EBSS (without phenol red) supplemented with collagenase (300U/mL), hyaluronidase (100U/mL), 2% FBS, gentamycin (100μg/mL), 100U/mL Pen/Strep, and doxycycline (2μg/mL) at 37°C for 4 hours. Cells were then resuspended in Dispase II (5mg/mL) and DNase I (100μg/mL) and filtered before plating. Mammosphere cultures were grown in RPMI-1640 media supplemented with 2μg/mL doxycycline, B27 (Invitrogen 17504-044), 10ng/mL murine EGF (Sigma E4127), 20ng/mL bFGF (Invitrogen 13256-029), Pen/Strep (Gibco 15140-122) and 2mM L-Glutamine (Gibco 25030-081). For microscopy experiments in Figure 1D and S2A, EGF and bFGF were excluded from the culture media. Mammospheres were passaged using enzymatic dissociation to re-plate as single cell suspensions. Doxycycline was excluded from the media where specified.

Primary and recurrent tumor cell lines were generated as previously described (Alvarez et al., 2013; Moody et al., 2005). Primary tumor cells were cultured in DMEM with 10% super calf serum, 1% Penicillin/Streptomycin, and 1% L-Glutamine supplemented with 10 ng/ml EGF, 5μg/ml insulin, 1μg/ml hydrocortisone, 5μg/ml prolactin, 1μM progesterone and 2μg/ml doxycycline to maintain HER2/neu expression. Recurrent tumor cells were cultured in DMEM with 10% SCS, 1% Penicillin/Streptomycin, and 1% L-Glutamine supplemented with 10 ng/ml EGF and 5 μg/ml insulin. Human Her2-amplified cells were obtained from the Duke University Cell Culture Facility, and sub-cultured per ATCC protocols.

### Metabolite Extraction

For mammosphere metabolomics experiment, 3x10^5^ cells for +dox condition and 6x10^5^ cells for -dox condition were plated in poly-hema coated 6-well plates. Mammospheres were collected by brief centrifugation (2 min, 300g), media were aspirated, and pellets were immediately resuspended in 1mL -80°C extraction solvent (80%MeOH/water). Tubes were moved to -80°C for 15 minutes before centrifugation (20,000g for 10 min at 4°C). Supernatant was transferred to a new tube, and samples were dried using a SpeedVac. Samples were stored at -80°C prior to LC-MS analysis. For primary and recurrent tumor cell adherent cultures, 3.5x10^5^ cells were plated in 6-well culture dishes. 24 hours later, media were quickly aspirated, and plates were placed on dry ice. 1mL -80°C extraction solvent (80%MeOH/water) was added to each well, and plates were moved to -80°C for 15 min. Plates were then scraped and collected in Eppendorf tubes for centrifugation (20,000g for 10 min at 4°C). Supernatant was transferred to a new tube, and samples were dried using a SpeedVac. Samples were stored at - 80°C prior to LC-MS analysis.

### Liquid Chromatography and Mass Spectrometry

Metabolite profiling was performed as previously described (Liberti et al., 2017; Liu et al., 2014). Ultimate 3000 UHPLC (Dionex) is coupled to Q Exactive Plus-Mass spectrometer (QE-MS, Thermo Scientific) for metabolite profiling. A hydrophilic interaction chromatography method (HILIC) employing an Xbridge amide column (100 x 2.1 mm i.d., 3.5 μm; Waters) is used for polar metabolite separation. Detailed LC method was described previously(Liu et al., 2014) except that mobile phase A was replaced with water containing 5 mM ammonium acetate (pH 6.8). The QE-MS is equipped with a HESI probe with related parameters set as below: heater temperature, 120 °C; sheath gas, 30; auxiliary gas, 10; sweep gas, 3; spray voltage, 3.0 kV for the positive mode and 2.5 kV for the negative mode; capillary temperature, 320 °C; S-lens, 55; A scan range (m/z) of 70 to 900 was used in positive mode from 1.31 to 12.5 minutes. For negative mode, a scan range of 70 to 900 was used from 1.31 to 6.6 minutes and then 100 to 1,000 from 6.61 to 12.5 minutes; resolution: 70000; automated gain control (AGC), 3 × 106 ions. Customized mass calibration was performed before data acquisition.

### Metabolomic Data Analysis

LC-MS peak extraction and integration were performed using commercial available software Sieve 2.2 (Thermo Scientific). The peak area was used to represent the relative abundance of each metabolite in different samples. The missing values were handled as described previously(Liu et al., 2014). The metabolite heatmap and pathway analysis were performed using MetaboAnalyst 4.0. For the heatmap, top 70 metabolites are displayed. For pathways analysis, all metabolites that had a 2-fold increase or decrease between recurrent cells expressing control or NRF2 targeting hairpin were analyzed.

### Cell viability assays

Unless otherwise indicated, cell viability assays were performed using CellTiter-Glo (Promega) according to manufacturer instructions. For adherent cell cultures, 1,000-5,000 cells were plated on an opaque 96-well plate. The next day, media were replaced with indicated drug treatments (see supplemental table for drug supplier and catalog numbers). For mammosphere cell viability experiment, cells were plated 2 days prior to drug treatment, and cells were collected by centrifugation and re-plated in media with or without dox and with or without 100μM etomoxir. The +dox samples were collected on day 3, and the -dox samples were collected on day 7. Media were refreshed once for the 7 day time point. Cells were collected by centrifugation and resuspended in CellTiter-Glo. They were then incubated at room temperature with shaking for 20 minutes before transferring to an opaque 96-well plate for luminescence measurement. For the SKBR3 Crystal Violet experiment, 1.25x10^5^ cells were plated on a 6-well plates. The next day, media were replaced with indicated media. After 6 days, media were removed, cells were rinsed once with cold PBS before incubating in crystal violet (0.5% crystal violet in 25% MeOH) for 5 min. Plates were rinsed in water 6 times before allowing to dry. For growth curve of recurrent tumor cells expressing scrambled or NRF2 targeting shRNA, 5x10^4^ cells were plated in 10cm dishes. Cells were counted the next day, and this was used as day 0. Cells were then counted on days 2, 4, and 6. Cells were counted using a hemocytometer.

### Immunoblotting and qPCR

Western blotting was performed as previously described (Alvarez et al., 2013). Primary and secondary antibodies used in this study are shown in Supplemental Table 1. For fluorescent antibodies membranes were imaged using Odyssey infrared imaging system (Li-Cor), and for HRP conjugated antibodies, membranes were treated with ECL reagents (Bio-Rad or Millipore Sigma) developed using autoradiography film (Genesee Scientific). Band intensities were quantified using ImageJ. RNA was extracted using the RNeasy kit (Qiagen). cDNA was synthesized using the ImProm-II Reverse Transcription System (Promega). Taqman probes used for qPCR are listed in Supplemental Table 1, and qPCR was performed using a CFX384 Touch Real-Time PCR Detection System (Bio-Rad). Actb and Gapdh were used for normalization controls. Reagent concentrations and supplier information are included in the Reagent table.

### DCFDA and mitoSOX assay

For DCFDA staining, cells were trypsinized and incubated in phenol red-free media containing 10% serum and 10μM DCFDA (Abcam) on a rocker at 37°C in the dark for 45 minutes. Cells were then collected by centrifugation, rinsed once with cold PBS, and resuspended in PBS supplemented with 0.1% serum. For MitoSOX staining, cells were incubated in phenol red-free media containing 10% serum and 2.5μM mitoSOX Red (Thermo) at 37°C for 10 minutes. Cells were then trypsinized, rinsed once with cold PBS, and resuspended in PBS supplemented with 0.1% serum. Both DCFDA and mitoSOX samples were immediately analyzed using a BD FACSCanto II flow cytometry. When indicated, samples were also imaged using Zeiss Axio Imager Widefield Fluorescence Microscope.

### Plasmids and viral transduction

To generate a lentiviral construct for expression of caNRF2, NRF2 (Nfe2l2) was PCR amplified from recurrent tumor cell cDNA. The primers (For-acttgtcgacgccaccatggattacaaagacgatgacgataaggcccagcacatccagacagacaccagtgg; Rev-atcagcggccgcactagtttttctttgtatctggcttcttgc), generated a truncated gene fragment lacking the first 88 codons, and included a kozak sequence, start codon, and flag epitope tag sequence. This was then cloned into the gateway entry vector pEntr4 (Thermo, A10465) using restriction digest (SalI and NotI). This was cloned into the gateway destination vector, pLenti PGK Neo DEST (w531-1), (a gift from Eric Campeau & Paul Kaufman; Addgene plasmid # 19067) (Campeau et al., 2009). The empty vector control was created by SalI restriction digest of the pLenti PGK Neo DEST (w531-1) destination vector to remove the ccdb cassette. The backbone was gel extracting and re-ligated. NRF2 targeting pLKO shRNA expression were obtained from Sigma (shNRF2#1: TRCN0000054659 and shNRF2#2: TRCN0000054658). Scramble shRNA was a gift from David Sabatini (Addgene plasmid # 1864) (Sarbassov et al., 2005).

To generate lentivirus, HEK293T cells were transfected with psPAX2 and pMDG.2 packaging plasmids (gifts from Didier Trono, EPFL, Lausanne, Switzerland; Addgene plasmids 12559 and 12660) and the lentiviral expression construct. Viral supernatant was collected after 48 and 72 hours and filtered. This virus was used to transduce cells with 6μg/mL polybrene (MilliporeSigma).

### Immunofluorescence

Mammospheres were collected by centrifugation onto coverslips. Frozen tissues were cut in 8μm sections (Figure 1) or 12μm sections (Figure 3) using a cryostat. Tissue sections or mammospheres were fixed in 4% paraformaldehyde, permeabilized in 0.5% Triton-X 100, and blocked in 3% BSA and 10% normal goat serum. Slides were incubated with primary antibodies for 1 hour at room temperature, washed, and incubated in with secondary AlexaFluor conjugated antibodies for 1 hour at room temperature. For lipid droplet imaging, cells were incubation with BODIPY for 30 min at room temperature before mounting coverslips with Prolong Gold. For experiments using Hoechst, Hoechst was included in the secondary antibody incubation. For experiments using DAPI, cells were mounted using Prolong Gold with DAPI. Slides were imaged using a Lecia SP5 inverted microscope. NRF2 nuclear intensity was quantified using CellProfiler (Broad Institute). Reagent concentrations and supplier information are included in the Reagent table.

### Human survival analysis

A publicly available online tool for correlating gene sets with survival in breast cancer was used. Gene expression-based Outcome for Breast cancer Online (GOBO (Ringner et al., 2011)) uses 4 datasets (GSE1456, GSE3494, GSE6532, and GSE7390), and it is available at http://co.bmc.lu.se/gobo/gsa.pl. Both the “NRF2 core gene signature” (previously compiled (Romero et al., 2017)) and the “NRF2 18-gene signature” were used for input. These gene sets are included in the Gene Signature table.

### Gene Set Enrichment Analysis (GSEA)

A publicly available RNA sequencing data of matched primary and metastatic tumors (GSE110590 (Siegel et al., 2018)) was downloaded. Expression values for metastases were averaged for each individual patient, and the difference (delta) between the primary tumor value and average metastasis value was calculated for each gene. Any genes with no value for the primary tumor were not included for that patient. A rank ordered list of the deltas for each individual patient was created, and the “NRF2 core gene signature” was used for GSEA. This gene set is included in the Gene Signature table. The differences for all individual patients were averaged (Average delta), and a rank ordered list was generated. The “NRF2 core gene signature” and “Hallmarks” gene signatures (Liberzon et al., 2015) were used for GSEA.

## Acknowledgements

We thank So Young Kim from the Duke Functional Genomics Core. We also thank members of the Alvarez lab for critical reading of this manuscript. This work was funded by National Cancer Institute grants R01CA208042 (JVA) and R01CA193256 (JWL), the V-Foundation, Golfers Against Cancer, the Integrative metabolomics shared resource, and by startup funds from the Duke Cancer Institute, the Duke University School of Medicine and the Whitehead Foundation (to JVA).

## Author Contributions

J.V.A. and D.B.F. were responsible for the conception, design, and interpretation of all experiments. M.D.H assisted in the conception and interpretation of metabolic experiments. D.B.F., R.L., L.C.N., and R.N. performed experiments and collected data. J.L. and J.W.L. designed, performed, and analyzed the metabolomics experiments. D.B.F. and J.V.A. wrote the manuscript. J.V.A. supervised all work.

