## Supplemental Figures and Tables for "NRF2-dependent metabolic reprogramming is required for tumor recurrence following oncogene inhibition"

**Supplemental Figure Legends**

**Figure S1:**

(A) Western blots for p-Her2 (Tyr1221/1222), p-ERK1/2 (Thr202/Tyr204), p-Akt (Ser473), and p-S6-RP (Ser240/244) in mammospheres cultured in the presence of dox (Her2 on) or without dox (Her2 off).

(B) Western blots for Her2, ERK1/2, Akt, and PCNA in mammospheres cultured in the presence of dox (Her2 on) or without dox (Her2 off).

(C) Western blot for p-PDH (Ser293 in mammospheres cultured in the presence of dox (Her2 on) or without dox (Her2 off) for 4 and 14 days.

**Figure S2:**

(A) DCFDA staining showing ROS levels in mammospheres cultured with dox or without dox for 2 days. Significance was determined by Student's t test (** $p<0.01$), and data are represented as mean ± SEM (n=6 fields of view).

(B) Relative viability of HCC1954 cells treated with lapatinib and 5 mM NAC for 72 hours. Significance was determined by two-way ANOVA (Tukey's multiple comparisons test; **$p<0.01$)

(C) DCFDA staining showing ROS levels in BT474 cells and SKBR3 cells treated with lapatinib or 100 µM DHEA for 48 hours. Significance was determined by one-way ANOVA with Tukey's multiple comparisons test (n=2).

(D) DCFDA staining showing ROS levels in BT474 cells treated with lapatinib and 50 µM 6-An for 48 hours. Significance was determined by two-way ANOVA with Tukey's multiple comparisons test (n=2).

(E) DCFDA staining showing ROS levels in mammospheres treated with 50 µM 6-An or TBHP for 24 hours. Significance was determined by Student's t test (n=2).

(F) Relative viability of mammosphere cultured with dox or without dox and with or without 100 µM etomoxir (eto) for 7 days. Significance was determined by two-way ANOVA with Tukey's multiple comparisons test (n=3).

(G) qPCR analysis of Cpt1a, Cpt1b, and CD36 expression in 2 independent mammosphere cultures grown in the presence of dox or without dox for 4 days. Significance was determined by two-way ANOVA with Tukey's multiple comparisons test (n=3).

(H) BODIPY staining showing lipid droplets (green) in mammospheres cultured in the presence of dox. Scale bar, 25 µm.

Error bars denote mean ± SEM. ns $p > 0.05$; *$p < 0.05$; **$p < 0.01$; ***$p < 0.001$.

**Figure S3:**

(A) qPCR analysis of Gclm, Nqo1, and Hmox1 expression in primary tumors (n=5) and tumors 48 hours after dox withdrawal (Her2 off; n=4). Significance was determined by two-way ANOVA with Tukey's multiple comparisons test.
(B) Immunofluorescence staining for NRF2 in a representative primary tumor (representative of 3 tumors) generated from MTB;TAN mice generated on 2mg/mL dox water and a residual tumor (representative of 2 tumors) generated from MTB;TAN mice on 0.1mg/mL dox water for primary tumor formation before 56 days without dox. Scale bar, 50µm.
(C) Waterfall plot of RNA sequencing data showing the expression of NRF2 target genes between primary tumor cell lines (n=2) and recurrent tumor cell lines (n=2).
Error bars denote mean ± SEM. **p < 0.01; ***p < 0.001.

**Figure S4:**
(A) DCFDA staining showing ROS levels in primary tumor cells expressing an empty vector (Cont) or caNRF2. Significance was determined by Student's t test (n=2).
(B) Kaplan-Meier plot showing primary tumor-free survival for mice bearing control or caNRF2 tumors. p-value was determined by log-rank (Mantel-Cox) test.
(C) Western blot for NRF2 and NQO1 in injected cells, primary tumors, and recurrent tumors expressing caNRF2.
(D) Kaplan-Meier plot showing recurrence-free survival (RFS) for breast cancer patients whose tumors have low (gray) or high (red) expression of the NRF2 core gene signature.
(E) Gene set enrichment analysis of matched primary and recurrent breast tumors showing enrichment of "Reactive Oxygen Species Pathway" and "Fatty Acid Oxidation" signatures in recurrent tumors.
(F) Expression of NRF2, KEAP1, TBP, and 10 canonical NRF2 target genes in matched primary and recurrent tumors from individual patients.
Error bars denote mean ± SEM. ns p > 0.05; **p < 0.01.

**Figure S5:**
(A) DCFDA staining showing ROS levels in two independent recurrent cell lines treated with 5 mM NAC.
Error bars denote mean ± SEM.

**Figure S6:**
(A) qPCR analysis of Taldo1 and Tkt expression in control (shScr) and NRF2-knockdown (shNRF2) recurrent tumor cells. Significance was determined by two-way ANOVA with Tukey's multiple comparisons test (n=2).

(B) Dose response curves showing relative viability in two primary and two recurrent tumor cell lines treated with indicated doses of BPTES for 48 hours.

(C) Dose response curves showing relative viability in control (shScr) and NRF2-knockdown (shNRF2) recurrent tumor cells treated with indicated doses of BPTES for 72 hours.

(D) Dose response curves showing relative viability in primary tumor cells expressing an empty vector (control) or constitutively active NRF2 (caNRF2) treated with indicated doses of BPTES for 72 hours.

(E) DCFDA staining showing ROS levels in control (shScr) and NRF2-knockdown (shNRF2) recurrent cells treated with 3 μM BPTES for 48 hours. Significance was determined by two-way ANOVA with Tukey's multiple comparisons test (n=2).

(F) Relative viability of recurrent cell lines treated with CB-839 and dimethyl-α-ketoglutarate (dm-αKG), as indicated. Significance was determined by two-way ANOVA with Tukey's multiple comparisons test (n=3).

Figure S1

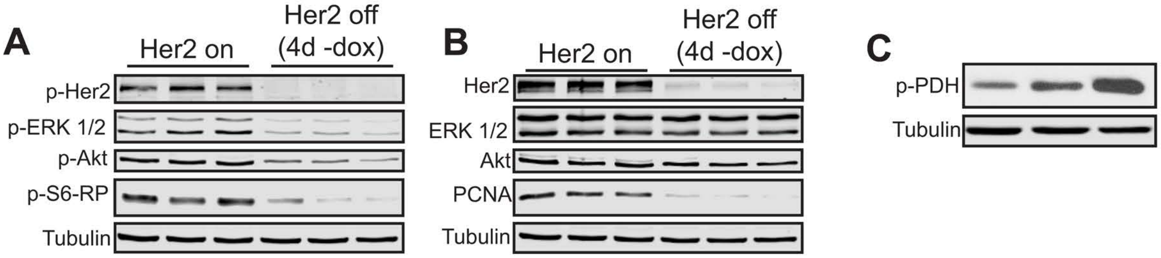
A
Her2 on
Her2 off
(4d -dox)
p-Her2
p-ERK 1/2
p-Akt
p-S6-RP
Tubulin
B
Her2 on
Her2 off
(4d -dox)
Her2
ERK 1/2
Akt
PCNA
Tubulin
C
p-PDH
Tubulin

Figure S2

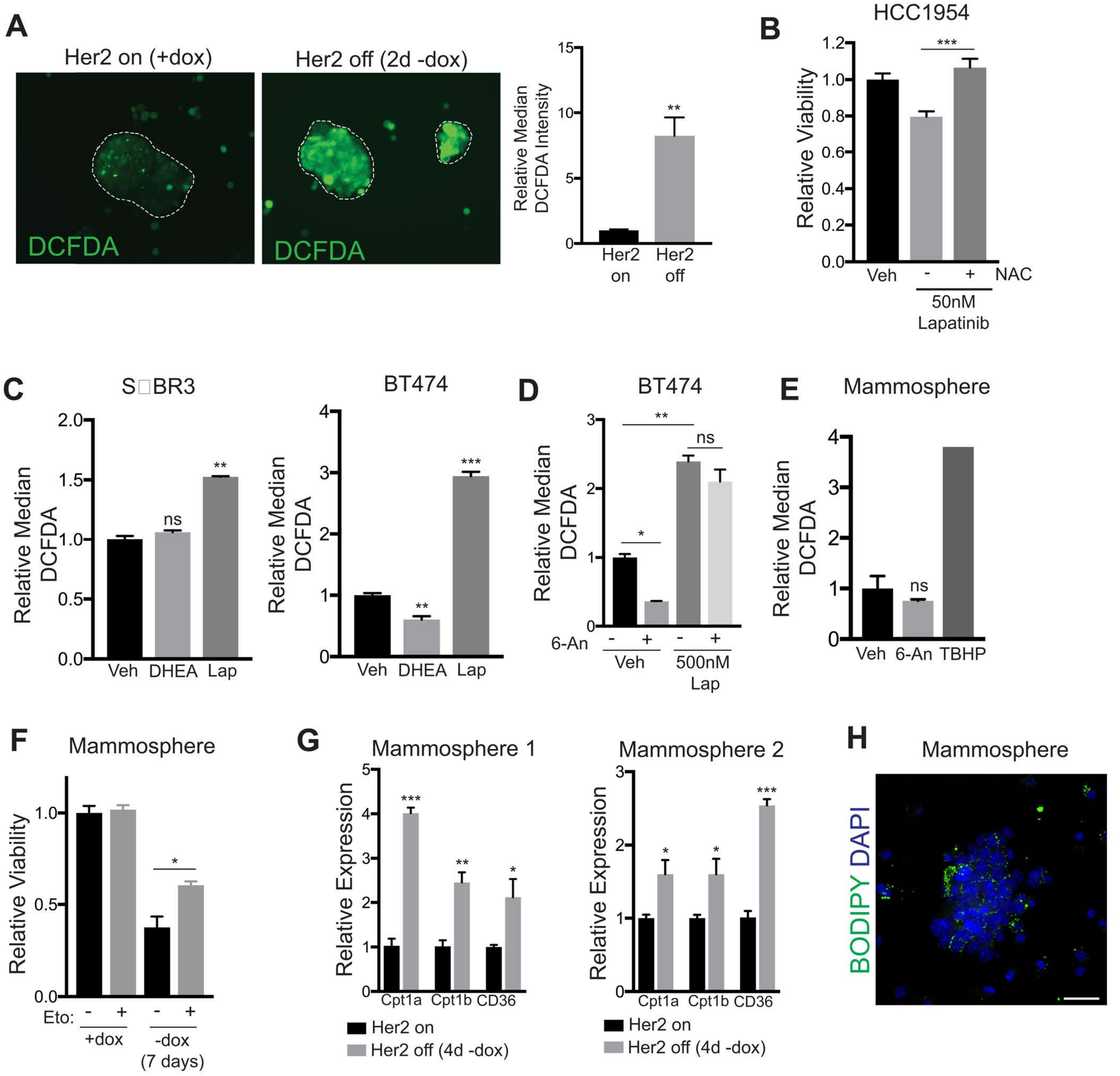
A
Her2 on (+dox)
Her2 off (2d -dox)
DCFDA
DCFDA
Relative Median DCFDA Intensity
15
10
5
0
**
Her2 on
Her2 off
B
HCC1954
Relative Viability
1.2
1.0
0.8
0.6
0.4
0.2
0.0
***
Veh
-
+
NAC
50nM Lapatinib
C
S□BR3
Relative Median DCFDA
2.0
1.5
1.0
0.5
0.0
ns
**
Veh
DHEA
Lap
BT474
Relative Median DCFDA
4
3
2
1
0
**
***
Veh
DHEA
Lap
D
BT474
Relative Median DCFDA
3
2
1
0
**
ns
*
6-An
-
+
-
+
Veh
500nM Lap
E
Mammosphere
Relative Median DCFDA
4
3
2
1
0
ns
Veh
6-An
TBHP
F
Mammosphere
Relative Viability
1.0
0.5
0.0
*
Eto:
-
+
-
+
+dox
-dox (7 days)
G
Mammosphere 1
Relative Expression
5
4
3
2
1
0
***
**
*
Cpt1a
Cpt1b
CD36
Her2 on
Her2 off (4d -dox)
Mammosphere 2
Relative Expression
3
2
1
0
*
*
***
Cpt1a
Cpt1b
CD36
Her2 on
Her2 off (4d -dox)
H
Mammosphere
BODIPY DAPI

Figure S3

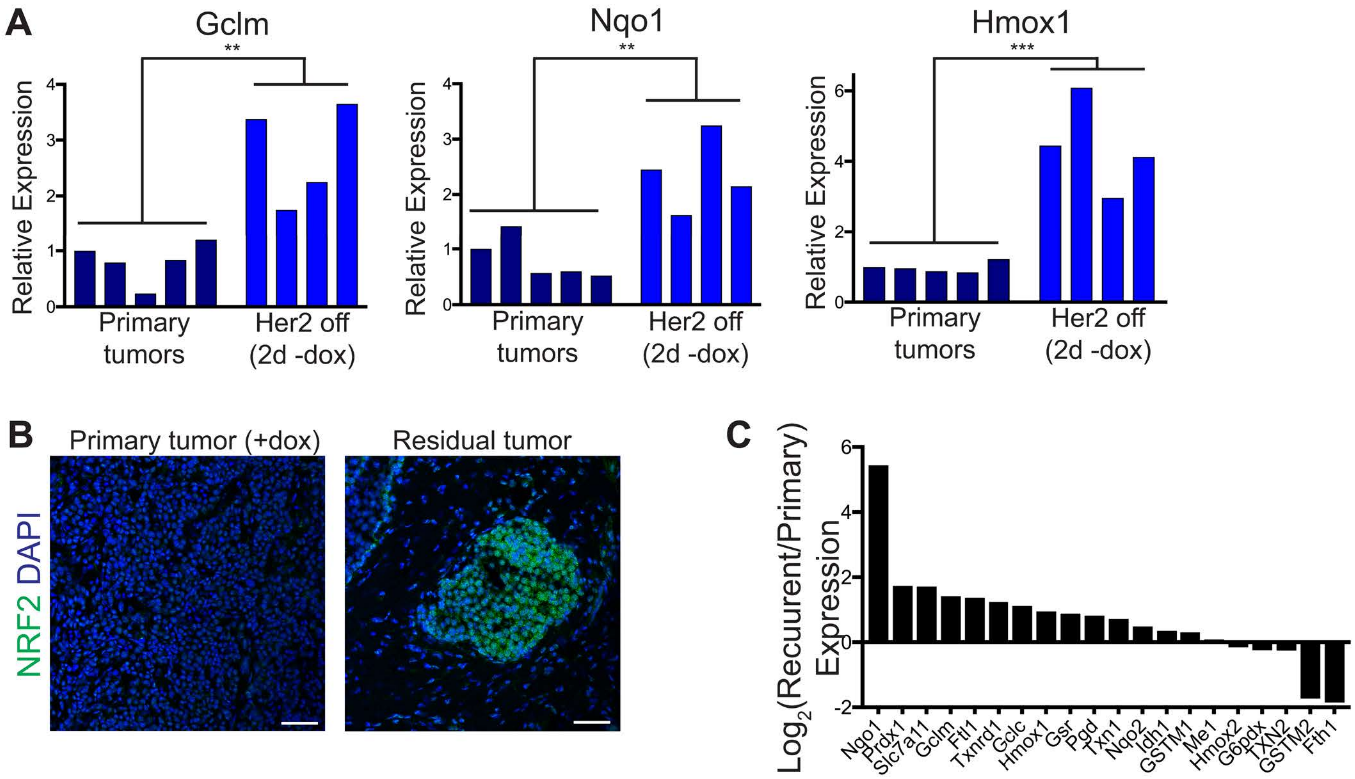
A
Gclm
**
Relative Expression
Primary tumors
Her2 off (2d -dox)
Nqo1
**
Hmox1
***
B
Primary tumor (+dox)
Residual tumor
NRF2 DAPI
C
Log2(Recuurent/Primary) Expression
Nqo1
Prdx1
Slc7a11
Gclm
Ftl1
Txnrd1
Gclc
Hmox1
Gsr
Pgd
Txn1
Nqo2
Idh1
GSTM1
Me1
Hmox2
G6pdx
TXN2
GSTM2
Fth1

Figure S4

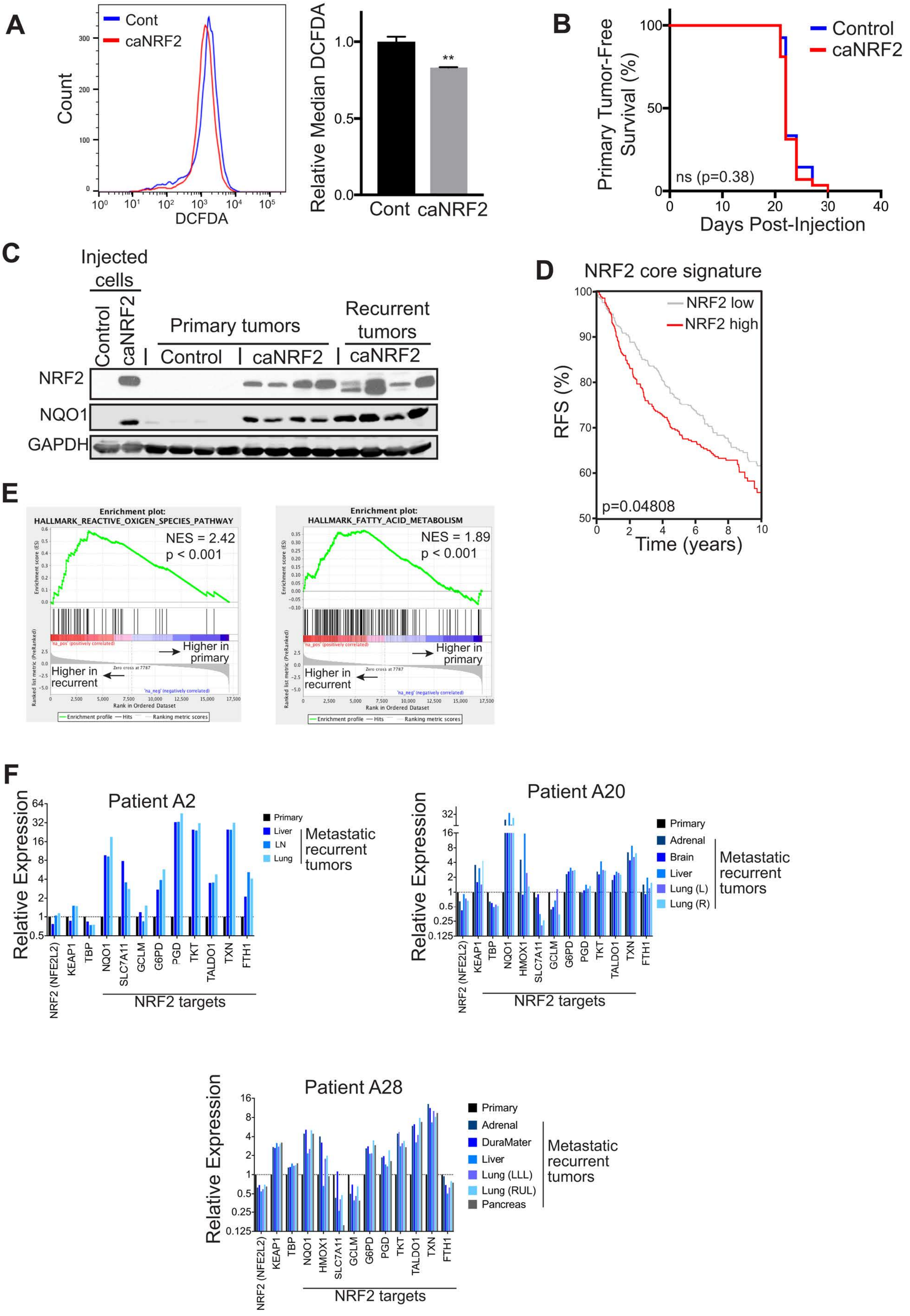
A
Cont
caNRF2
Count
DCFDA
Relative Median DCFDA
**
Cont
caNRF2
B
Control
caNRF2
Primary Tumor-Free Survival (%)
ns (p=0.38)
Days Post-Injection
C
Injected cells
Control
caNRF2
Primary tumors
Recurrent tumors
Control
caNRF2
caNRF2
NRF2
NQO1
GAPDH
D
NRF2 core signature
NRF2 low
NRF2 high
RFS (%)
p=0.04808
Time (years)
E
Enrichment plot:
HALLMARK_REACTIVE_OXIGEN_SPECIES_PATHWAY
NES = 2.42
p < 0.001
Higher in primary
Higher in recurrent
Enrichment plot:
HALLMARK_FATTY_ACID_METABOLISM
NES = 1.89
p < 0.001
Higher in primary
Higher in recurrent
F
Patient A2
Relative Expression
Primary
Liver
LN
Lung
Metastatic recurrent tumors
NRF2 (NFE2L2)
KEAP1
TBP
NQO1
SLC7A11
GCLM
G6PD
PGD
TKT
TALDO1
TXN
FTH1
NRF2 targets
Patient A20
Relative Expression
Primary
Adrenal
Brain
Liver
Lung (L)
Lung (R)
Metastatic recurrent tumors
NRF2 (NFE2L2)
KEAP1
TBP
NQO1
HMOX1
SLC7A11
GCLM
G6PD
PGD
TKT
TALDO1
TXN
FTH1
NRF2 targets
Patient A28
Relative Expression
Primary
Adrenal
DuraMater
Liver
Lung (LLL)
Lung (RUL)
Pancreas
Metastatic recurrent tumors
NRF2 (NFE2L2)
KEAP1
TBP
NQO1
HMOX1
SLC7A11
GCLM
G6PD
PGD
TKT
TALDO1
TXN
FTH1
NRF2 targets

Figure S5

A

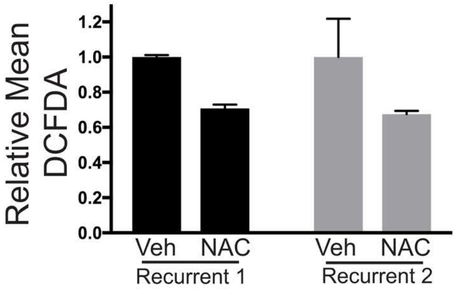
Relative Mean
DCFDA
1.2
1.0
0.8
0.6
0.4
0.2
0.0
Veh
NAC
Recurrent 1
Veh
NAC
Recurrent 2

Figure S6

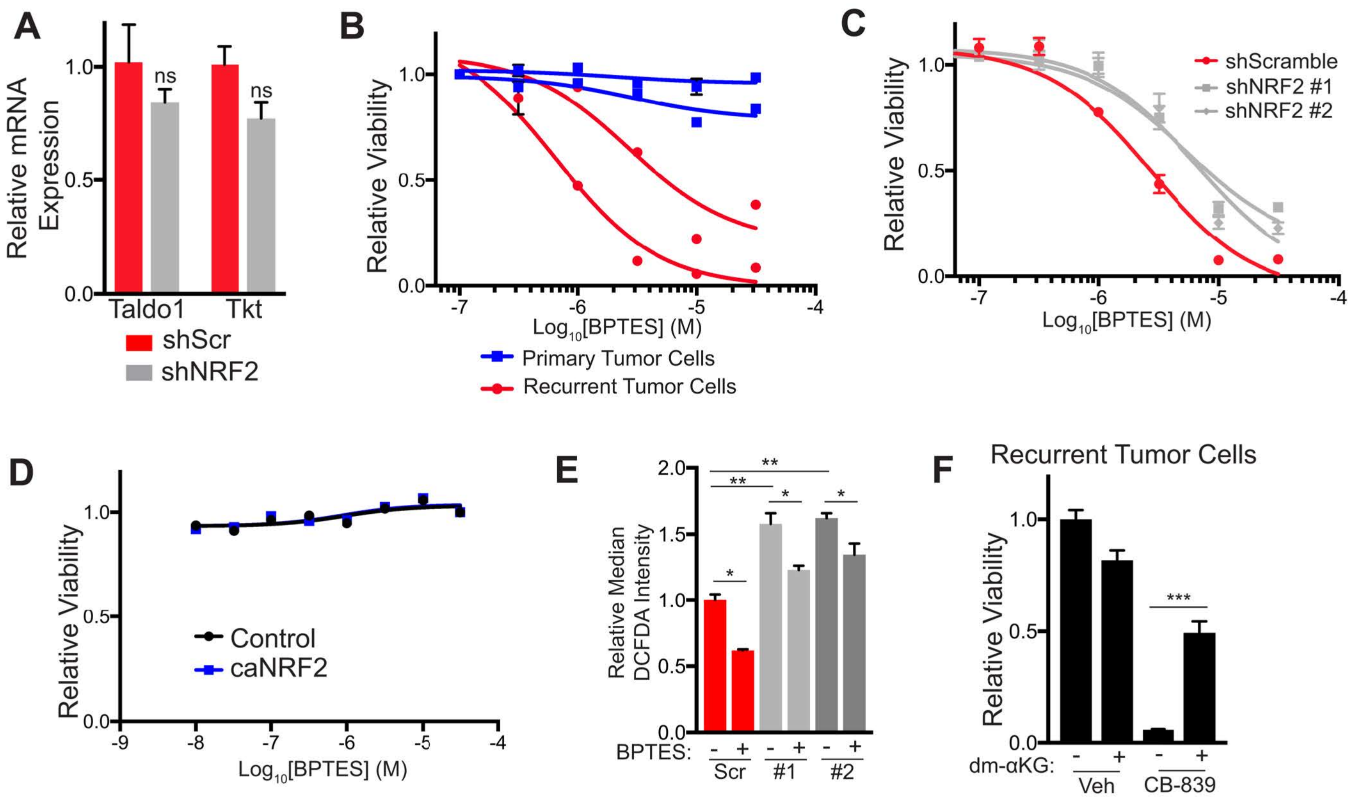
A
Relative mRNA Expression
1.0
0.5
0.0
ns
ns
Taldo1
Tkt
shScr
shNRF2
B
Relative Viability
1.0
0.5
0.0
-7
-6
-5
-4
Log10[BPTES] (M)
Primary Tumor Cells
Recurrent Tumor Cells
C
Relative Viability
1.0
0.5
0.0
-7
-6
-5
-4
Log10[BPTES] (M)
shScramble
shNRF2 #1
shNRF2 #2
D
Relative Viability
1.0
0.5
0.0
-9
-8
-7
-6
-5
-4
Log10[BPTES] (M)
Control
caNRF2
E
Relative Median DCFDA Intensity
2.0
1.5
1.0
0.5
0.0
**
**
*
*
*
BPTES:
-
+
-
+
-
+
Scr
#1
#2
F
Recurrent Tumor Cells
Relative Viability
1.0
0.5
0.0
***
dm-αKG:
-
+
-
+
Veh
CB-839

| NRF2 Core Gene Signature | |
|---|---|
| Gene name | Notes |
| ABCB6 | |
| ABCC1 | |
| ABCC2 | |
| ABCG2 | |
| ABHD4 | |
| ADAM23 | |
| AKR1B10 | |
| AKR1C1 | |
| AKR1C3 | |
| ALDH3A1 | |
| ALDH3A2 | |
| ASF1A | |
| ASNS | |
| ASPH | |
| BLVRB | |
| C14ORF149 | 1 |
| C16ORF28 | 1,2 |
| C1S | |
| CAMKK1 | 1 |
| CBR1 | |
| CBR3 | |
| CCDC77 | 1 |
| CCND3 | |
| CDKN2B | |
| CES1 | |
| CLN5 | |
| CPLX2 | |
| CXCR7 | |
| CYP4F11 | |
| DDC | |
| DEGS1 | |
| DGKG | |
| DKFZP762E1312 | 1,2 |
| DOCK10 | |
| EPHX1 | |
| F2RL2 | |
| FAM55C | |
| FTH1 | |
| FZD7 | |
| G6PD | |
| GALNT13 | 1 |
| GCLC | |
| GCLM | |
| GLA | |
| GPC1 | |
| GPX2 | |
| GSR | |
| HGD | |
| HMOX1 | |
| HRB | 1,2 |
| HSPA1B | |
| HTATIP2 | |
| IDH1 | |
| KIAA0319 | |
| LOC642252 | 1,2 |
| LOC644799 | 1,2 |
| LRP12 | |
| LRP8 | |
| LTB4DH | 1,2 |
| MAFG | |
| MAP2 | |
| MARS | |
| MCM10 | |

| NRF2 18-gene Signature |
|---|
| Gene name |
| FTH1 |
| G6PD |
| GCLC |
| GCLM |
| GPX2 |
| GSR |
| HMOX1 |
| HMOX2 |
| IDH1 |
| ME1 |
| NQO1 |
| NQO2 |
| PGD |
| PRDX1 |
| SLC7A11 |
| TXN |
| TXN2 |
| TXNRD1 |

| | |
|---|---|
| ME1 | |
| MEGF9 | |
| MGT1 | 1 |
| MSC | |
| NCF2 | |
| NEIL3 | |
| NFE2L2 | |
| NLN | 1 |
| NQO1 | |
| NQO2 | |
| NR0B1 | 2 |
| NRCAM | |
| OSGIN1 | |
| P2RY6 | |
| PA2G4 | |
| PBEF1 | 1,2 |
| PCK2 | |
| PGD | |
| PIR | |
| POPDC3 | |
| PPAT | |
| PRDX1 | |
| RRM2 | |
| SERPINE1 | |
| SFN | |
| SLC38A6 | |
| SLC6A6 | |
| SLC7A11 | |
| SORD | |
| SPANXA1 | 1,2 |
| SPANXB1 | 1,2 |
| SPP1 | |
| SQSTM1 | |
| SRXN1 | 1 |
| TALDO1 | |
| TFRC | |
| TGFB2 | |
| TKT | |
| TM4SF20 | 2 |
| TRIM16 | |
| TSPAN7 | |
| TXN | |
| TXNRD1 | |
| UCHL1 | |
| UIP1 | 1,2 |

From Romero, R. et al. Keap1 loss promotes Kras-driven lung cancer and results in dependence on glutaminolysis. Nat Med 23, 1362-1368 (2017).
1: Gene excluded from survival analysis (not found in human gene expression datasets).
2: Gene excluded from GSEA (not found in human recurrent tumor gene expression datasets).

Reagents

| Antibodies | | | | | |
|---|---|---|---|---|---|
| **Protein** | **Application** | **Supplier** | **Catalog #** | **Concentration** | **Additional notes** |
| Cleaved PARP | WB | Cell Signaling | 9544 | 1:1000 | |
| p-AMPK (Thr172) | WB | Cell Signaling | 2535 | 1:1000 | |
| AMPKα | WB | Cell Signaling | 5831 | 1:1000 | |
| GAPDH | WB | Sigma | G8795 | 1:5000 | |
| p-Her2 (Tyr1221/1222) | WB | Cell Signaling | 2243 | 1:1000 | |
| Her2 | WB | Cell Signaling | 4290 | 1:1000 | |
| p-ERK1/2 (Thr202/Tyr204) | WB | Cell Signaling | 9106 | 1:2000 | |
| ERK1/2 | WB | Cell Signaling | 9102 | 1:1000 | |
| p-Akt (Ser473) | WB | Cell Signaling | 4060 | 1:1000 | |
| Akt | WB | Cell Signaling | 9272 | 1:1000 | |
| p-S6-RP (Ser240/244) | WB | Cell Signaling | 5364 | 1:2000 | |
| α-Tubulin | WB | Santa Cruz | 8035 | 1:2000 | Used for all except Figs. S1C, 4B, 5B, 6G |
| α-Tubulin | WB | Cell Signaling | 3873 | 1:5000 | Used for Figs. S1C, 4B, 5B, 6G |
| p-PDH | WB | Calbiochem | AP10062 | 1:10000 | |
| NRF2 | WB | Cell Signaling | 12721 | 1:1000 | |
| NQO1 | WB | Cell Signaling | 62262 | 1:1000 | |
| KEAP1 | WB | Proteintech | 10503-2-AP | 1:1000 | |
| Her2 | IF | Cell Signaling | 2165 | 1:200 | |
| Cleaved Caspase-3 (Asp175) | IF | Cell Signaling | 9661 | 1:300 | |
| Ki67 - AlexaFluor 488 | IF | BD Pharmingen | 561165 | 1:100 | |
| NRF2 | IF | GeneTex | GTX103322 | 1:250 | |
| Annexin V - AlexaFluor 488 | Flow | Life Technologies | V13245 | 1:20 | |
| Goat anti-rabbit AlexaFluor 488 | IF | Life Technologies | A11034 | 1:5000 | |
| Goat anti-rabbit AlexaFluor 568 | IF | Life Technologies | A11036 | 1:5000 | |
| Goat anti-rabbit AlexaFluor 680 | WB | Life Technologies | A21076 | 1:5000 | |
| Goat anti-rabbit IRDye® 800CW | WB | LiCor | 925-32211 | 1:5000 | |
| Goat anti-mouse IRDye® 800CW | WB | LiCor | 926-32210 | 1:5000 | |
| Anti-rabbit IgG, HRP-linked antibody | WB | Cell Signaling | 7074P2 | 1:5000 | |

| Other dyes and staining materials | | | | | |
|---|---|---|---|---|---|
| **Compound** | **Application** | **Supplier** | **Catalog #** | **Concentration** | **Additional Notes** |
| BODIPY 493/503 | IF | Thermo | D3922 | | |
| Hoechst 33342 | IF | Thermo | H1399 | 2µg/mL | |
| DCFDA | IF, Flow | Abcam | ab113851 | 10µM | |
| mitoSOX Red | Flow | Thermo | M36008 | 2.5µM | |
| Prolong Gold with DAPI | IF | Cell Signaling | 8961 | | Used for all images with DAPI |
| Prolong Gold | IF | Thermo | P36930 | | Used for all images with Hoechst |

| Taqman gene expression probes | | | |
|---|---|---|---|
| Taqman Master mix | Applied Biosystems #4369016 | | |
| **Gene** | **Species** | **Catalog #** | **Additional notes** |
| Actb | mouse | Mm02619580_g1 | |
| Gapdh | mouse | Mm99999915_g1 | Used for Fig 6E |
| Cpt1a | mouse | Mm01231183_m1 | |
| Cpt1b | mouse | Mm00487191_g1 | |
| Gclm | mouse | Mm01324400_m1 | |
| Nqo1 | mouse | Mm01253561_m1 | |
| Hmox1 | mouse | Mm00516005_m1 | |
| Slc7a11 | mouse | Mm00442530_m1 | |
| NRF2 (Nfe2l2) | mouse | Mm00477784_m1 | |
| G6pdx | mouse | Mm00656735_g1 | |
| Pgd | mouse | Mm00503037_m1 | |
| Taldo1 | mouse | Mm00807080_g1 | |
| Tkt | mouse | Mm00447559_m1 | |

| SYBR green gene expression primers | | | |
|---|---|---|---|
| SYBR green master mix | Bio-Rad #172-5121 | | |
| **Gene** | **Forward sequence** | **Reverse Sequence** | **Additional notes** |
| CD36 | GAACCACTGCTTTCAAAAACTGG | TGCTGTTCTTTGCCACGTCA | |
| Actb | CATGAAGATCCTGACCGAGCGTG | TCTGCTGGAAGGTGGACAGTGAGG | |
| Nqo1 | TATCCTTCCGAGTCATCTCTAGCA | TCTGCAGCTTCCAGCTTCTTG | Used for Fig S3A |
| Hmox1 | CAGGTGATGCTGACAGAGGA | GAGAGTGAGGACCCACTGGA | Used for Fig S3A |
| Gclm | GCCACCAGATTTGACTGCCTTTG | TGCTCTTCACGATGACCGAGTACC | Used for Fig S3A |

| Cell culture reagents | | | | |
|---|---|---|---|---|
| **Compound** | **Supplier** | **Catalog #** | **Concentration** | **Additional notes** |
| RPMI-1640 | Sigma | R8758 | | |
| DMEM | Corning | 10-017-CV | | |
| Pen/Strep | Gibco | 15140-122 | 1x | |
| L-Glutamine | Gibco | 25030-081 | 2mM | |
| Fetal Bovine Serum (FBS) | Corning | 35-010-CV | | |
| Super Calf Serum (SCS) | Gemini | 100-510 | | |
| EGF | Sigma | E4127 | 10ng/mL | |
| bFGF | Invitrogen | 13256-029 | 20ng/mL | |
| B27 Supplement | Invitrogen | 17504-044 | 1x | |
| Poly(2-hydroxyethyl methacrylate) | Sigma | P3932 | | |
| Doxycycline | RPI | D43020 | 2µg/mL | |

| | | | | |
|---|---|---|---|---|
| Insulin | Sigma | I6634 | 5µg/mL | |
| Prolactin | National hormone and peptide program | | 5µg/mL | Provided by A. F. Parlow (Harbor-UCLA Medical Center) |
| hydrocortisone | Sigma | H0396 | 1µg/mL | |
| Progesterone | Sigma | P7556 | 1µM | |

| **Cell treatment compounds** | | | |
|---|---|---|---|
| **Compound** | **Supplier** | **catalog #** | **Additional Notes** |
| Lapatinib | Selleckchem | S1028 | |
| N-acetyl cysteine (NAC) | Sigma | A9165 | |
| Glutathione | Sigma | G6013 | |
| Etomoxir | Tocris | 4539 | |
| 6-Aminonicotinamide | Cayman Chemical | 10009315 | |
| DHEA | Sigma | D4000 | |
| TBHP | Abcam | ab113851 | From DCFDA kit |
| TBHQ | Sigma | 112941 | |
| BPTES | Sigma | SML0601-5MG | |
| CB-839 | Cayman | 22038 | |
| dimethyl α-ketoglutarate | Sigma | 349631 | |
